# Structural Insight into Mitochondrial β-Barrel Outer Membrane Protein Biogenesis

**DOI:** 10.1101/2020.04.09.034678

**Authors:** Kathryn A. Diederichs, Xiaodan Ni, Sarah E. Rollauer, Istvan Botos, Xiaofeng Tan, Martin S. King, Edmund R.S. Kunji, Jiansen Jiang, Susan K. Buchanan

**Affiliations:** Laboratory of Molecular Biology, National Institute of Diabetes & Digestive & Kidney Diseases, National Institutes of Health, 9000 Rockville Pike, Bethesda, MD 20892 USA; Laboratory of Membrane Protein Structural Biology, National Heart, Lung, and Blood Institute, National Institutes of Health, Bethesda, MD 20892 USA; MRC Mitochondrial Biology Unit, University of Cambridge, Cambridge Biomedical Campus, Cambridge CB2 0XY, UK

## Abstract

In mitochondria, β-barrel outer membrane proteins mediate protein import, metabolite transport, lipid transport, and biogenesis. The Sorting and Assembly Machinery (SAM) complex consists of three proteins that assemble as a 1:1:1 complex to fold β-barrel proteins and insert them into the mitochondrial outer membrane. We report cryoEM structures of the SAM complex from *Myceliophthora thermophila*, which show that Sam50 forms a 16-stranded transmembrane β-barrel with a single polypeptide-transport-associated (POTRA) domain extending into the intermembrane space. Sam35 and Sam37 are located on the cytosolic side of the outer membrane, with Sam35 capping Sam50, and Sam37 interacting extensively with Sam35. Sam35 and Sam37 each adopt a GST-like fold, with no functional, structural, or sequence similarity to their bacterial counterparts. Structural analysis shows how the Sam50 β-barrel opens a lateral gate to accommodate its substrates. The SAM complex structure suggests how it interacts with other mitochondrial outer membrane proteins to create supercomplexes.

## Introduction

β-barrel membrane proteins are found in the outer membranes of mitochondria, chloroplasts, and Gram-negative bacteria. Mitochondrial β-barrel membrane proteins are synthesized in the cytosol and imported across the mitochondrial outer membrane as unfolded precursor proteins by the Translocase of the Outer Membrane (TOM) complex. In the intermembrane space, β-barrel precursor proteins interact with small Translocase of the Inner Membrane (TIM) chaperones prior to their transfer to the Sorting and Assembly Machinery (SAM) complex for folding and insertion into the outer membrane^1^. The SAM complex consists of three components: a β-barrel core, Sam50, which spans the outer membrane, and two accessory subunits, Sam35 and Sam37, that associate with Sam50 on the cytosolic side of the membrane^2-5^. Sam50 and Sam35 are essential proteins, with Sam50 folding and inserting β-barrel substrates into the outer membrane and Sam35 interacting with the substrate β-signal located in the last β-strand^3^. Sam37, while not essential, functions in substrate release^3,6^ and may also promote formation of a TOM-SAM supercomplex^7^.

Bacterial β-barrel membrane proteins take a different pathway to reach the outer membrane, but they are inserted by evolutionarily related machinery^8^. In bacteria, proteins are synthesized in the cytoplasm, secreted across the inner membrane by the Sec translocon, bound to chaperones in the periplasm, and transferred to the Bacterial Assembly Machinery (BAM) complex for folding and insertion into the outer membrane^9,10^. In *Escherichia coli*, four lipoproteins, BamB, BamC, BamD, and BamE, associate with the periplasmic domain of BamA (itself a 16-stranded transmembrane β-barrel) to fold and insert β-barrels ranging in size and complexity from 8 β-strands with a simple barrel fold, to 26 β-strands with multiple domains^11^. Structures of BamA and BAM complexes illustrate two features of BamA that facilitate folding and insertion: [1] BamA has a narrowed hydrophobic surface where the first and last β-strands meet, which locally compresses and destabilizes the lipid bilayer to allow membrane protein insertion (BamA assisted model). [2] The first and last β-strands form a ‘lateral gate’ that has been shown to open and close by molecular dynamics (MD) simulations, disulfide crosslinking, and structures solved by X-ray crystallography and cryo-electron microscopy (cryoEM)^12-19^. The required opening and closing of the lateral gate gave rise to the budding model, where a β-hairpin (or larger portion) of the substrate enters the BamA lumen, associates with unpaired β-strands on BamA, and uses the BamA β-barrel to sequentially fold into the outer membrane, budding off from BamA when the first and last strands of the substrate come together. With the wide variety and large quantities of β-barrel proteins that exist in bacteria, it is possible that either or both models, or variations of these^20,21^, are used depending on the complexity of the substrate.

Sam50 and BamA are evolutionarily related, and both are members of the Omp85 superfamily^8,22^. Although no structures had previously been determined for any of the SAM components, a homology model based on BamA was used to study mitochondrial membrane protein insertion by crosslinking precursor proteins to predicted strands β1 and β16 of Sam50, suggesting that substrates enter the lumen of Sam50, accumulate at the lateral gate, and are released into the compressed membrane adjacent to the gate^23^. In contrast to bacteria, mitochondria are predicted to make only four types of β-barrel outer membrane protein: Sam50, Tom40, VDAC, and Mdm10^24^. Structures of Tom40^25,26^ and VDAC^27^ reveal very similar 19-stranded β-barrel proteins; Mdm10 is expected to adopt the same fold. Therefore, it appears that Sam50 substrates may use a single folding and insertion mechanism.

Although evolutionarily related, a number of important differences between SAM and BAM exist, suggesting that the details of how substrates are targeted to the folding machinery, where and how those substrates interact with peripheral SAM and BAM components, the functions of those peripheral components, and how substrates are released into the membrane once folded, will differ substantially. First, the entry pathways clearly differ as described above. Mitochondrial proteins enter from outside the organelle but interact with Sam50 on the intermembrane space side of the outer membrane. In contrast, bacterial substrates in the periplasm associate with periplasmic lipoproteins BamB, BamC, BamD, and BamE. BamD is essential in this process and has been shown to interact with unfolded substrates, activating BamA for substrate folding^28^. The peripheral SAM components, Sam35 and Sam37, sit on the opposite side of the membrane and therefore cannot make the same types of interactions with substrates that BAM lipoproteins do. Second, mitochondria contain only one POTRA domain while bacteria contain multiple POTRA domains^22^. In bacteria, the POTRA domain nearest the β-barrel is essential for activity^29^, while in mitochondria it can be removed without major consequence^3^. Third, mitochondrial Sam35 and Sam37 have been demonstrated to have functions not found in BAM lipoproteins^6^.

To better understand how β-barrel proteins are folded in mitochondria, we used cryoEM to solve structures of the complete SAM complex in lipid nanodiscs at 3.4 Å resolution, and in detergent to 3.0 Å resolution, with five structures in total (one in lipid, four in detergent). The SAM complex consists of one copy each of Sam50, which spans the mitochondrial outer membrane, and Sam35 and Sam37, which sit on the cytosolic side of the membrane. Sam50 forms a 16-stranded β-barrel which is minimally closed between strands β1 and β16. The single POTRA domain extends into the intermembrane space away from the barrel lumen, potentially allowing substrate access from this side of the membrane. The N-terminal portion of Sam35 interacts extensively with Sam50 on the cytosolic side of the membrane, occluding substrate efflux. Sam35 also interacts extensively with Sam37, such that Sam37 makes no contacts with Sam50 on the cytosolic side of the membrane; however, the linker between two predicted transmembrane *α*-helices in Sam37 interacts with the Sam50 POTRA domain in the intermembrane space. A comparison of the structures shows how Sam50 opens a lateral gate to fold and insert substrates, while an analysis of the SAM complex shows how it may interact with TOM to form TOM-SAM supercomplexes.

## Results

### Preparation of SAM complexes

To obtain a homogeneous SAM complex, we co-expressed *M. thermophila* (recently renamed to *Thermothelomyces thermophilus*) Sam50, Sam35, and Sam37 in *Saccharomyces cerevisiae* (see Methods for details). The SAM complex was solubilized from isolated mitochondria using the detergent lauryl maltose neopentyl glycol (LMNG) and purified by affinity chromatography using a Twin-Strep tag on the N-terminus of Sam37. Before incorporating into lipid nanodiscs, the SAM complex was further purified on a size exclusion column using LMNG. For single particle analysis in detergent, LMNG was exchanged on the size exclusion column for glycol-diosgenin (GDN), a synthetic substitute for digitonin. The purified samples contained approximately stoichiometric amounts of each of the three subunits.

### The SAM complex is monomeric in lipid nanodiscs

The cryoEM structure of the SAM complex in lipid nanodiscs was reconstructed from 179,509 particles and yielded a 3.4 Å resolution structure (Fig. 1, Supplementary Fig. 1 and Table 1). The resolution was sufficient to allow *ab initio* tracing of a majority of the folds of all three subunits. In lipid nanodiscs, the SAM complex is monomeric and contains one copy each of Sam50, Sam35, and Sam37. Sam50 forms a 16-stranded transmembrane β-barrel with a single POTRA domain extending into the intermembrane space. Sam35 and Sam37 are located on the cytosolic side of the membrane, where Sam35 caps the barrel lumen of Sam50 and also interacts extensively with Sam37 (Fig. 1B, C). The orientations of Sam35 and Sam37, as well as visualization of the lipid nanodisc, suggest that Sam35 and Sam37 interact peripherally with the outer leaflet of the membrane (Fig. 1A) and residues 339-353 of Sam37 form an amphipathic *α*-helix that appears to interact with the membrane (Fig. 1B). In this structure, the highest resolution is observed for the Sam50 β-barrel and for Sam35, with lower resolution (and more flexibility) observed for Sam37 and the Sam50 POTRA domain (Supplementary Fig. 1D). At low resolution, the first of two predicted transmembrane *α*-helices in Sam37 is visible; however, it is not visible at high resolution. It appears that the relatively large size of the lipid nanodisc compared to the Sam50 β-barrel prevents optimal particle alignment and further resolution improvement. Nonetheless, this is the first high resolution structure of an Omp85 family member solved in a lipid environment.

**Table 1.**
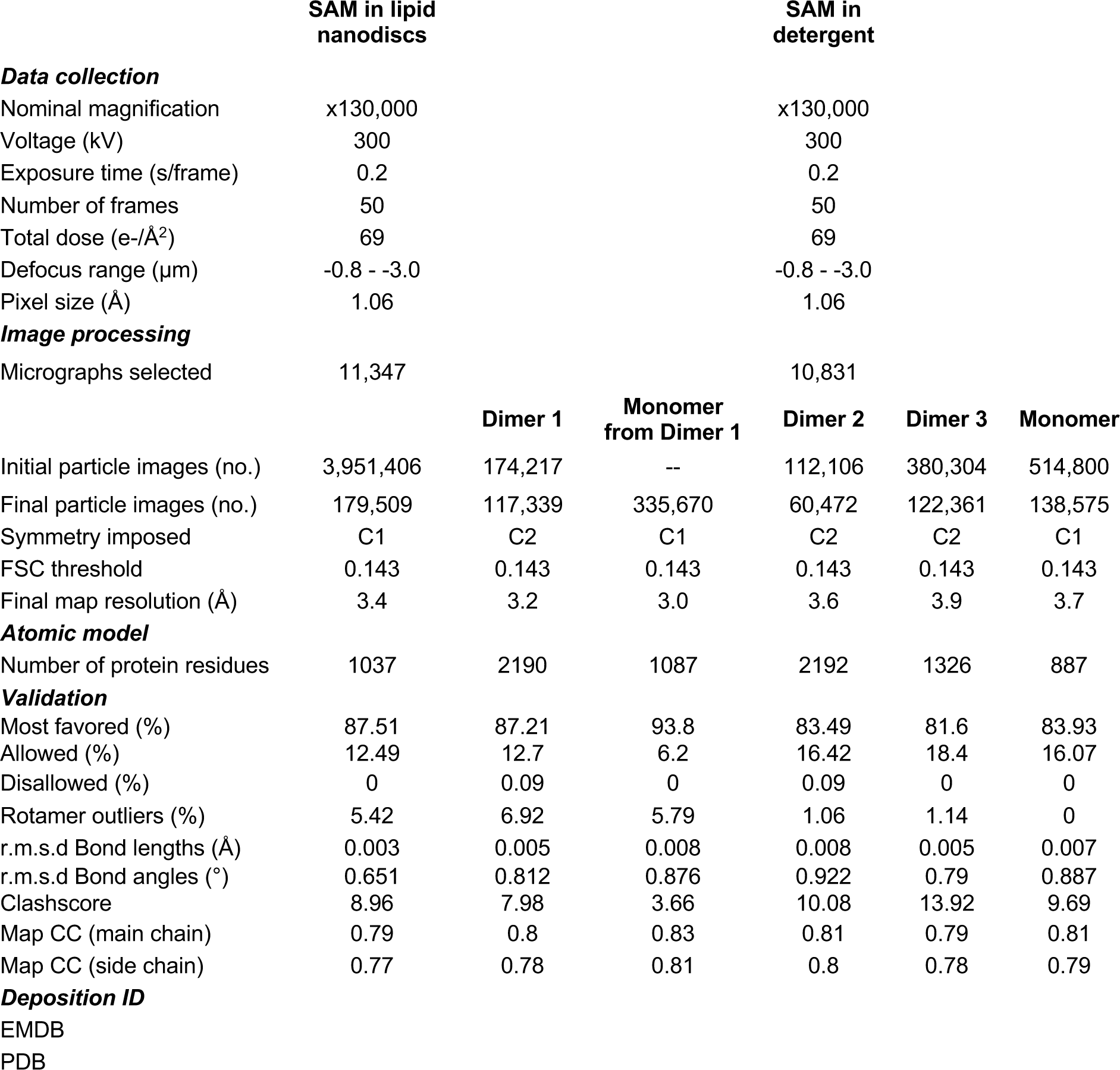
CryoEM data collection, structure determination and model statistics.

**Figure 1.**
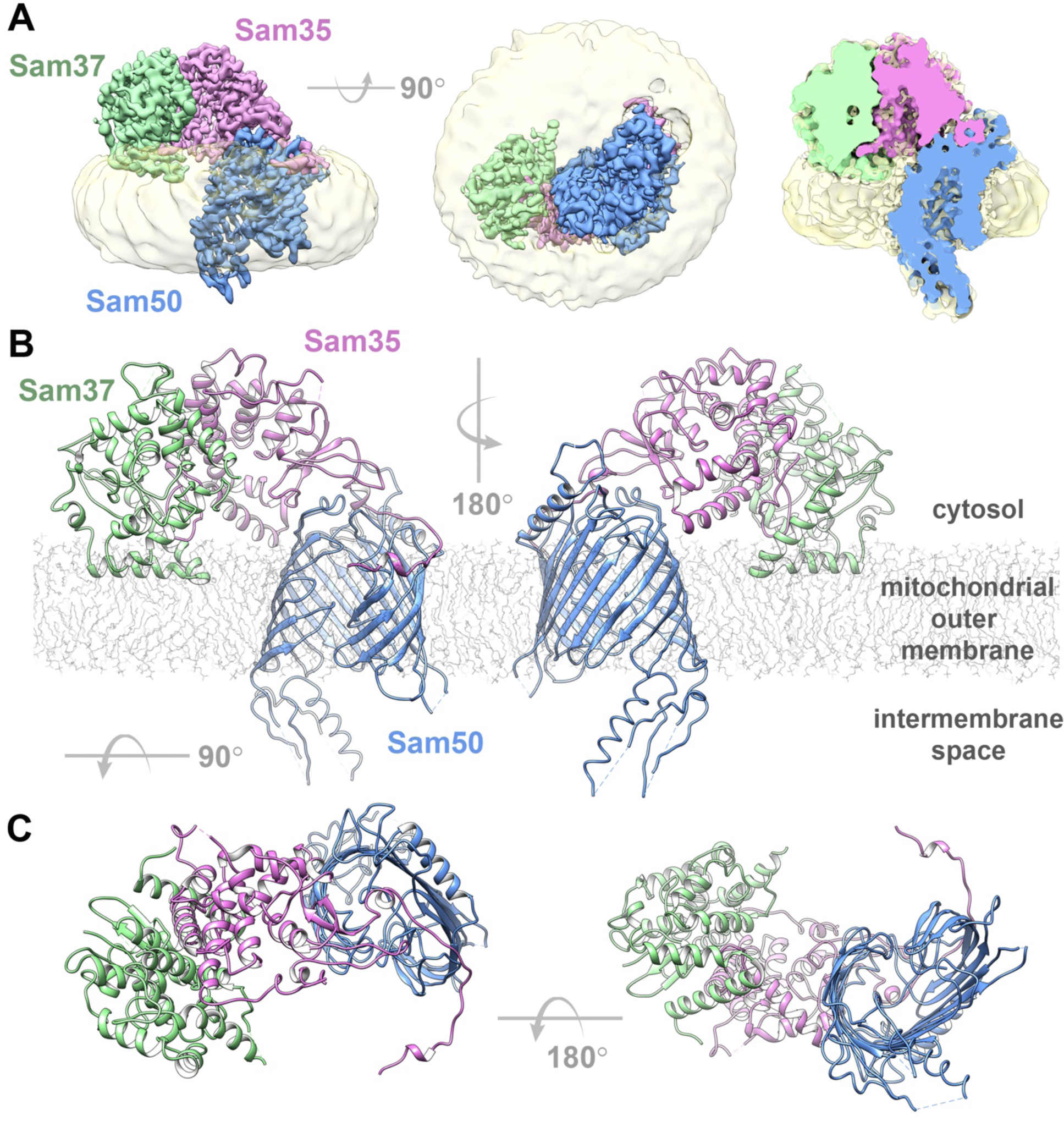
Structure of the SAM complex in lipid nanodiscs. (**A**) The cryoEM density map of the SAM complex (Sam35 in orchid, Sam37 in pale green, and Sam50 in blue) with the nanodisc densities (transparent yellow) showing the membrane boundaries. A cross section is shown on the right. (**B**) Ribbon views of the SAM complex in the context of a model lipid bilayer. (**C**) Ribbon views from top (cytosol) and bottom (intermembrane space). See also Supplementary Fig. 1 and Table 1.

### The SAM complex forms dimers in detergent

In addition to using lipid nanodiscs, we determined the SAM complex structure in the detergent GDN using cryoEM (see Methods for details). In this sample, 3D reconstructions exhibited several conformations, with monomer and multiple dimer conformations present (here, the term monomer refers to the SAM complex, with one copy each of Sam50, Sam35, and Sam37; the dimer refers to two SAM complexes) (Fig. 2, Supplementary Fig. 2, 3,4 and Table 1). The monomer conformation, although determined to a resolution of only 3.7 Å, is virtually identical to the monomer determined in lipid nanodiscs and to the monomer derived from dimer 1, described below. All of the dimers appear to be non-physiological, with an up-down association of SAM complexes not anticipated from functional studies. However, we were able to determine a structure of the most prevalent dimer (dimer 1) at 3.2 Å resolution from 117,339 particles (Fig. 2A). The SAM complex monomer determined from lipid nanodiscs was fitted into half of dimer 1, revealing that they are in an almost identical conformation (Fig. 2B). Therefore, we performed particle symmetry expansion in dimer 1, focused on only half of the dimer, and proceeded to refine this monomer to a final resolution of 3.0 Å, resulting in more clearly resolved sidechain densities (Fig. S2 and Supplementary Fig. S3). The first transmembrane *α*-helix of Sam37 is well defined, spanning the membrane adjacent to the Sam50 β-barrel, but making no contact with the β-barrel. This contrasts with the recently reported high resolution structures of the TOM core complex, where transmembrane *α*-helices closely associate with the hydrophobic surface of the Tom40 β-barrel^25,26^. The linker that connects the two predicted transmembrane helices in Sam37 interacts with the Sam50 POTRA domain, contributing an additional anti-parallel β-strand to that domain.

**Figure 2.**
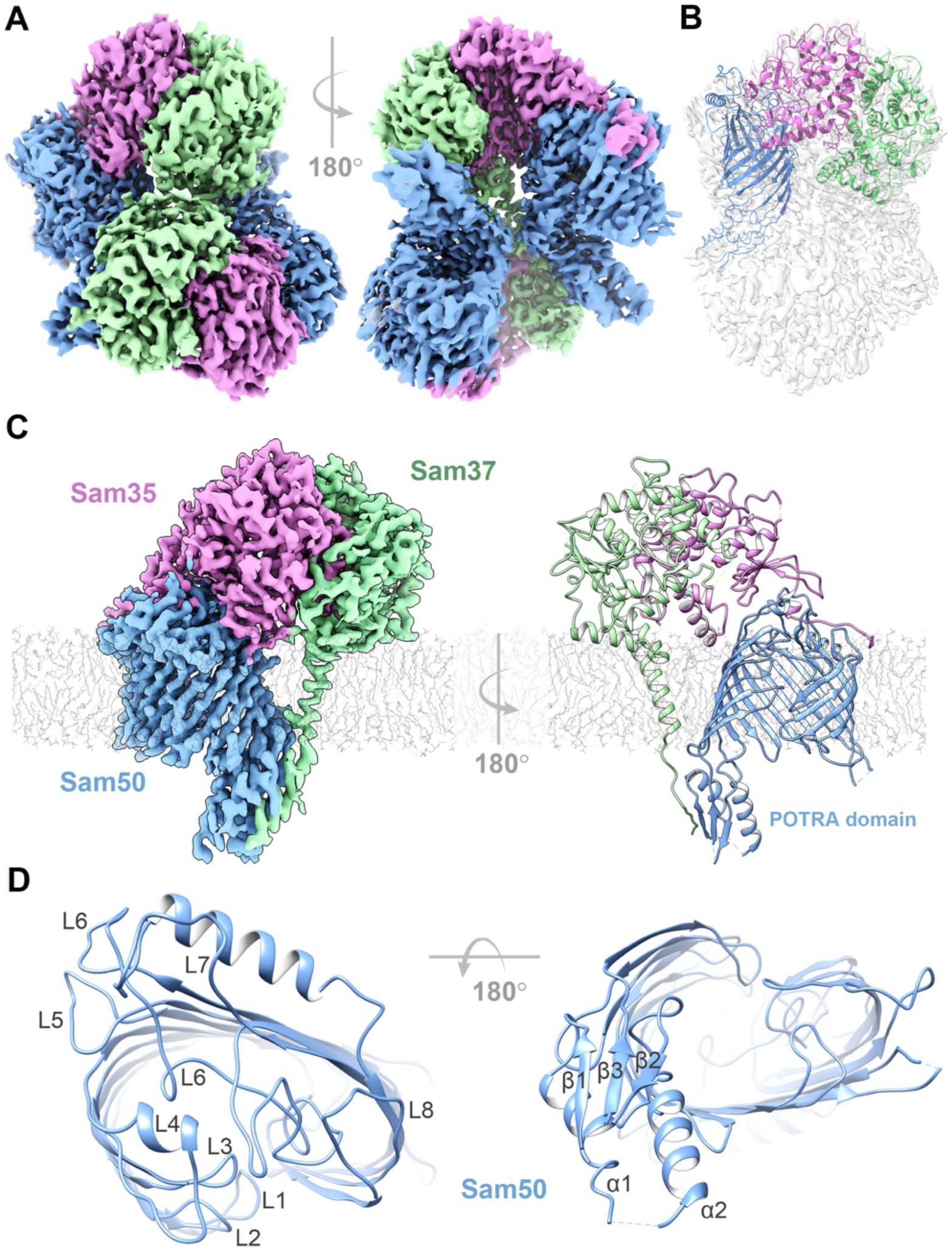
Structure of the SAM complex in detergent GDN. (**A**) The cryoEM density map of a dimer of the SAM complex (dimer 1) in GDN with Sam 35 in orchid, Sam37 in pale green and Sam50 in blue. (**B**) Superposition of the structure of the SAM complex in nanodiscs (ribbons) to the cryoEM density map of dimer 1 (transparent surface). (**C**) Side view of the high-resolution structure of the SAM complex reconstructed from the dimer 1 particles using symmetry expansion. The cryoEM density map and the atomic model are shown on the left and right, respectively. (**D**) The top and bottom views of Sam50 barrel showing the cytosolic loops L1-L8 and the POTRA domain with a βααββ fold. See also Supplementary Fig. 2, 3, 4, 5, and Table 1.

**Figure 3.**
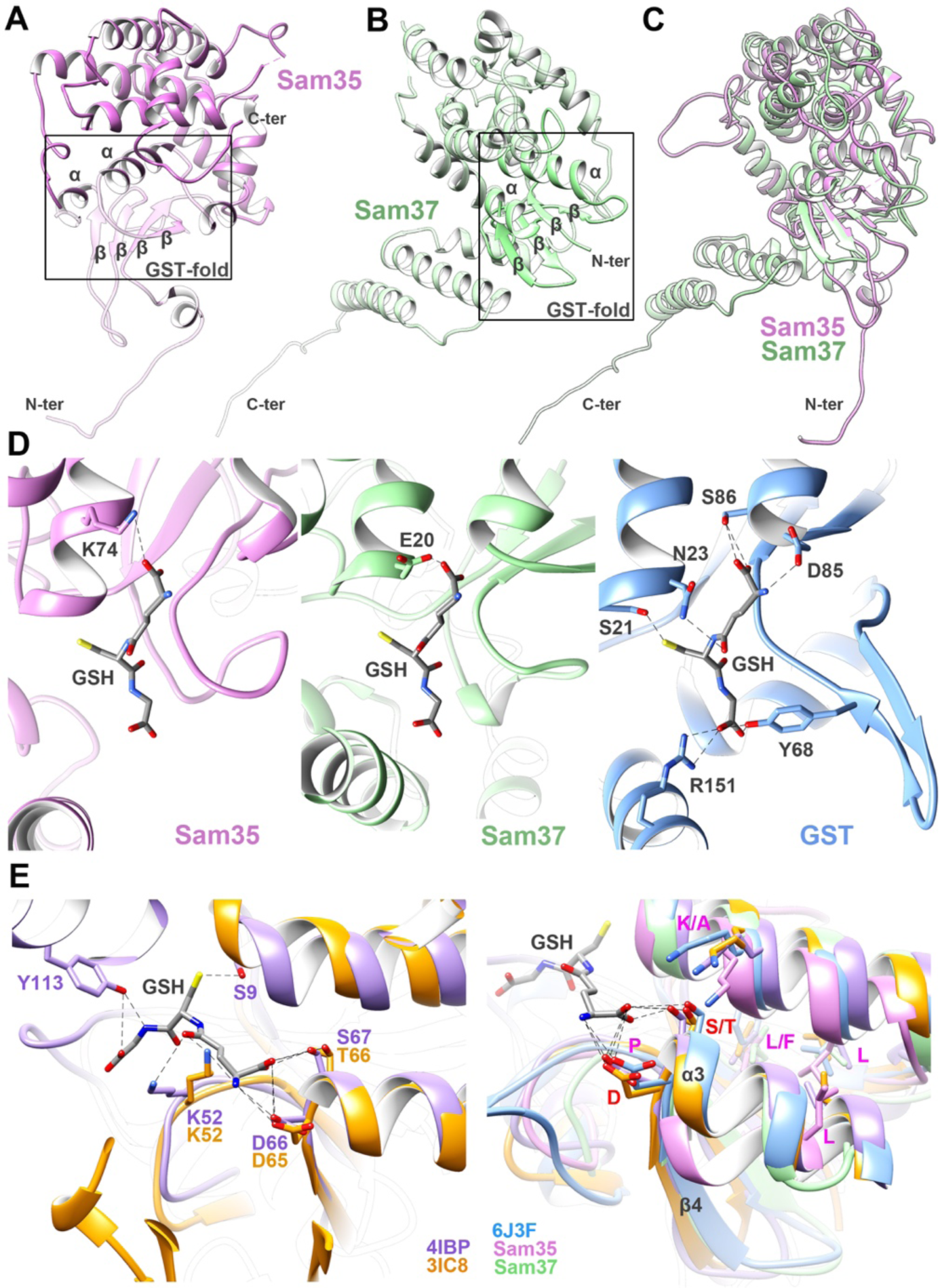
Analysis of the GST-like fold for Sam35 and Sam37. (**A**) Ribbon view of Sam35 exhibits a canonical GST fold of an α/β domain at the N-terminus and an all α-helical C-terminal domain. **(B**) Ribbon view of Sam37 shows a configuration similar to Sam35 with an N-terminal α/β domain with an all α-helical C-terminal domain. **(C)** Superposition of Sam35 (orchid) and Sam37 (pale green) shows high structural similarity. (**D**) Close-up of the pseudo active site in Sam35, Sam37, and the genuine active site of GST (PDB:6J3F). Hydrogen bonds are shown as dashed lines. (**E**) Close-up views of the residues related to the active site of GST. Left: superposition of two GST-like proteins (left; PDB:4IBP and PDB:3IC8). Right: superposition of GST (PDB:6J3F), the same two GST-like proteins (PDB:4IBP and PDB:3IC8), Sam35 (orchid), and Sam37 (pale green). The bound GSH is shown in grey. The active site residues conserved among all the above structures are labeled in magenta, whereas residues only present in GST and GST-like proteins are labeled in red.

**Figure 4.**
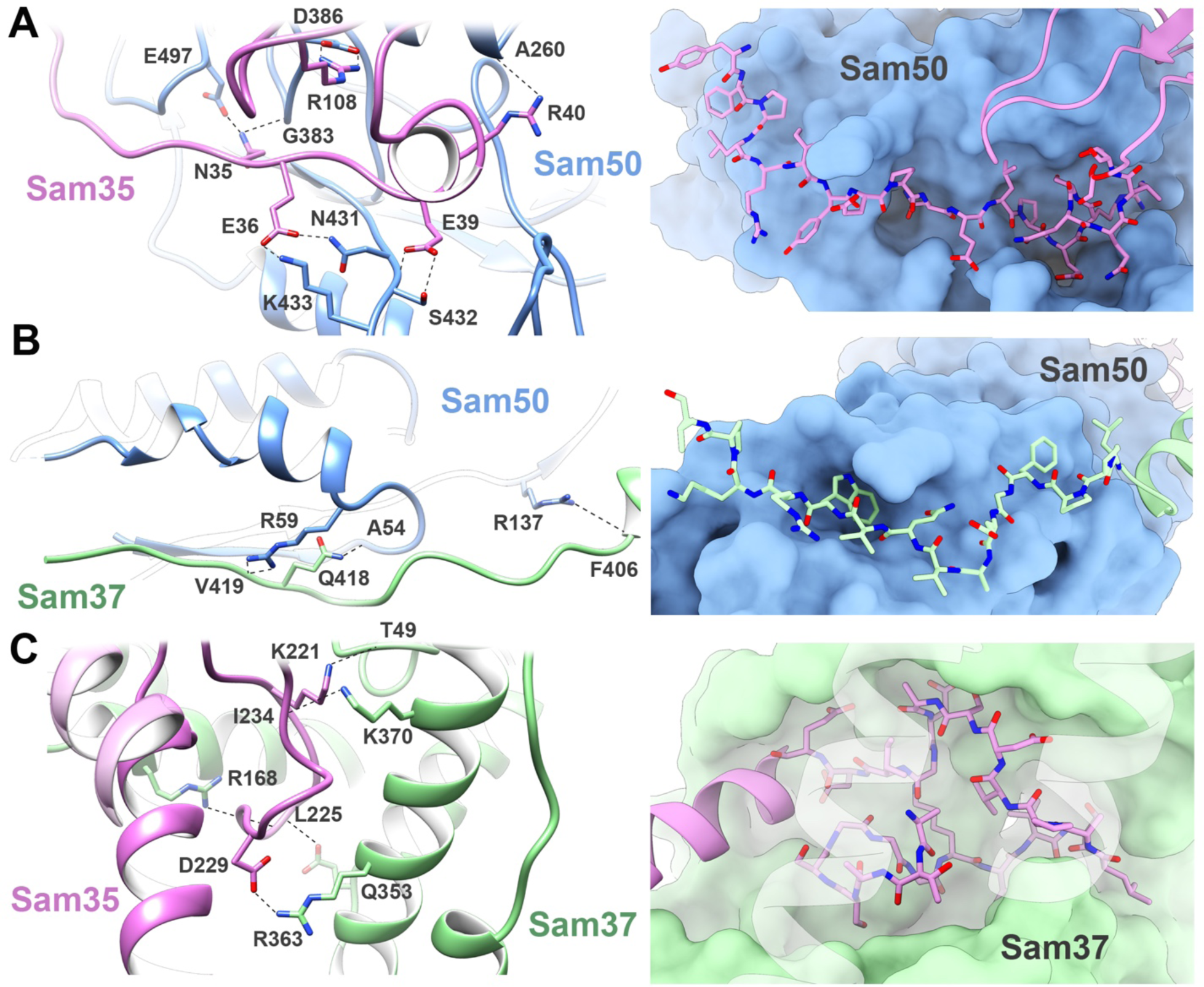
Interactions between individual components of the SAM complex. Interfaces for **(A)** Sam35/Sam50 β-barrel, **(B)** Sam50 POTRA/Sam37, and **(C)** Sam35/Sam37, in cartoon and surface representations. The interacting residues are labeled. See also Supplementary Fig. 6 and Supplementary Table 2, 3.

### The Sam50 β-barrel is only partially closed

As had been predicted from homology to BamA^19^, Sam50 consists of a 16-stranded transmembrane β-barrel preceded by a single POTRA domain in the intermembrane space (Fig. 2C, D and Supplementary Fig. 5). The POTRA domain adopts the classic *βααββ* fold, and is positioned away from the barrel lumen, allowing substrate access in this conformation. Structures of Omp85 proteins illustrate how mobile POTRA domains can be, potentially allowing or blocking access to the barrel lumen from the periplasm (bacteria) or intermembrane space (mitochondria). The reduced resolution in this domain in the lipid nanodisc structure further illustrates its mobility (Supplementary Fig. 1D). On the cytosolic side of the membrane, Sam50 loops L1 – L8 extend from the surface, positioning loops L4, L7, and L8 to interact with several N-terminal residues of Sam35. Loop 7 contains a surface-exposed *α*-helix that sits parallel to the membrane, while L6 extends deep into the empty barrel. These interactions effectively close the β-barrel on the cytosolic side, preventing substrate efflux. In both lipid and detergent, the Sam50 β-barrel adopts a kidney bean shape, with dimensions 50 by 40 Å (Fig. 1C, 2D and Supplementary Fig. 4). It is only partially closed at the interface, making no direct interactions between strands β1 and β16 (Supplementary Fig. 7). The short lengths of strands β1-β4 and β15-β16, and their orientation in the membrane, suggest that the lipid bilayer is likely to be compressed and disordered in this region, as has been observed for BamA^18,19^. We note that the conformation of the interface between strands β1 and β16 is virtually identical in lipid and detergent, allowing all subsequent analyses to be carried out with the higher resolution detergent-based structure of Sam50 (Supplementary Fig. 4D).

**Figure 5.**
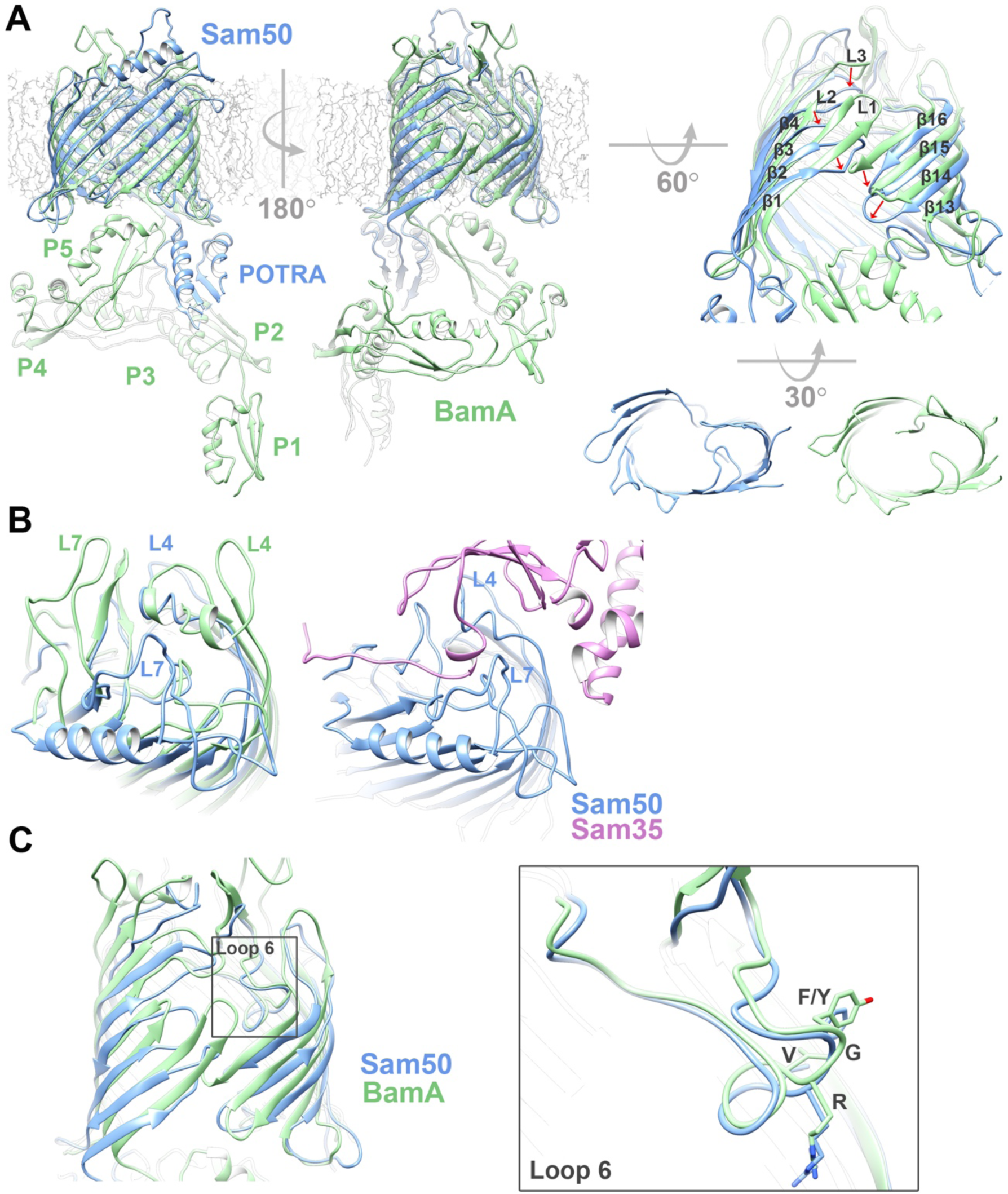
Comparison of Sam50 to BamA. (**A**) Superposition of Sam50 (blue) and BamA (green PDB:4K3B Right: close-up of the Sam50 lateral gate in detergent GDN compared to BamA and a bottom view of the barrel. The displacement of loops between the two structures is shown by red arrows. (**B**) Differences between Sam50 and BamA in outward facing loops L4 and L7 in the context of Sam35. (**C**) Superposition of loop 6 between the two structures. The conserved (V/I)RG(F/Y) motif is labeled. See also Supplementary Fig. 5, 7, Supplementary Table 4, 5 and Supplementary Movie 1.

### Sam35 and Sam37 adopt a GST-like fold

Sam35 and Sam37 share no sequence similarity with any of the BAM lipoproteins. The structures of Sam35 and Sam37 reveal that they each adopt a GST-like fold, which is also completely different from the BAM lipoproteins. Sam35 and Sam37 exhibit canonical N-terminal *α*/β domains and all *α*-helical C-terminal domains (Fig. 3A, B). Superposition of Sam35 and Sam37 shows the extensive structural similarities (Fig. 3C). Although Sam35 and Sam37 belong to the family of GST-like proteins, they do not have the necessary active site residues to catalyze conjugation of glutathione to a substrate (Fig. 3D). We compared the true GST protein (PDB:6J3F) with GST-like proteins (PDB:4IBP and PDB:3IC8), including Sam35 and Sam37 (Fig. 3E). Superpositions for Sam35 and GST, or for Sam37 and GST, have RMSDs of 2.5 Å and 2.2 Å, respectively. Clearly, secondary structure elements align well but active site residues are missing. Members of the GST protein family exhibit extreme diversity in sequence, and a large proportion of them are of unknown function. Sam35 plays an essential role in the SAM complex, and has been shown to bind precursor protein^3^. However, the SAM complex structure does not currently offer any clues to this interaction. Sam37 is not essential, but has been shown to function later in the folding process, facilitating precursor release^6^. It is interesting that the Sam37 linker binds to the POTRA domain of Sam50 in the intermembrane space, since the POTRA domain has also been implicated in precursor release^30^. This interaction places the two subunits with a shared function in close proximity. Our structure supports the previous observations that Sam37 serves to stabilize the interaction of Sam35 with Sam50, as there are extensive interactions between cytosolic Sam37 and Sam35^6,31^.

### Subunit interactions between Sam50, Sam35, and Sam37

Sam35 interacts with a groove in Sam50 created by loops L4 and L7, primarily through residues near its N-terminus. The interactions consist of 15 hydrogen bonds and 3 salt bridges resulting in a buried surface area of 3918.6 Å^2^ (Fig. 4A and Supplementary Tables 1, 2). These interactions allow Sam35 to sit tightly along the edge of the Sam50 β-barrel. In the intermembrane space, the Sam50 POTRA domain interacts with the linker connecting the two predicted transmembrane helices of Sam37 through 8 hydrogen bonds, resulting in a buried surface area of 2789.8 Å^2^ (Fig. 4B and Supplementary Tables 1, 2). Sam35 and Sam37 interact extensively on the cytosolic side of the membrane through 9 hydrogen bonds and 2 salt bridges, for a total buried surface area of 2978.5 Å^2^ (Fig. 4C and Supplementary Tables 1, 2).

Of the conserved residues from structure-based sequence alignments (Supplementary Fig. 5, 6 and Supplementary Table 3), only two interactions occur at the subunit interfaces involving highly conserved residues: Tyr156 in Sam35 with semi-conserved Asp111 on Sam37, and Gly383 in Sam50 with non-conserved Asn35 on Sam 35 (Supplementary Fig. 6 and Supplementary Table 3). Otherwise, there are no clear patterns of conserved residues in subunit interactions.

**Figure 6.**
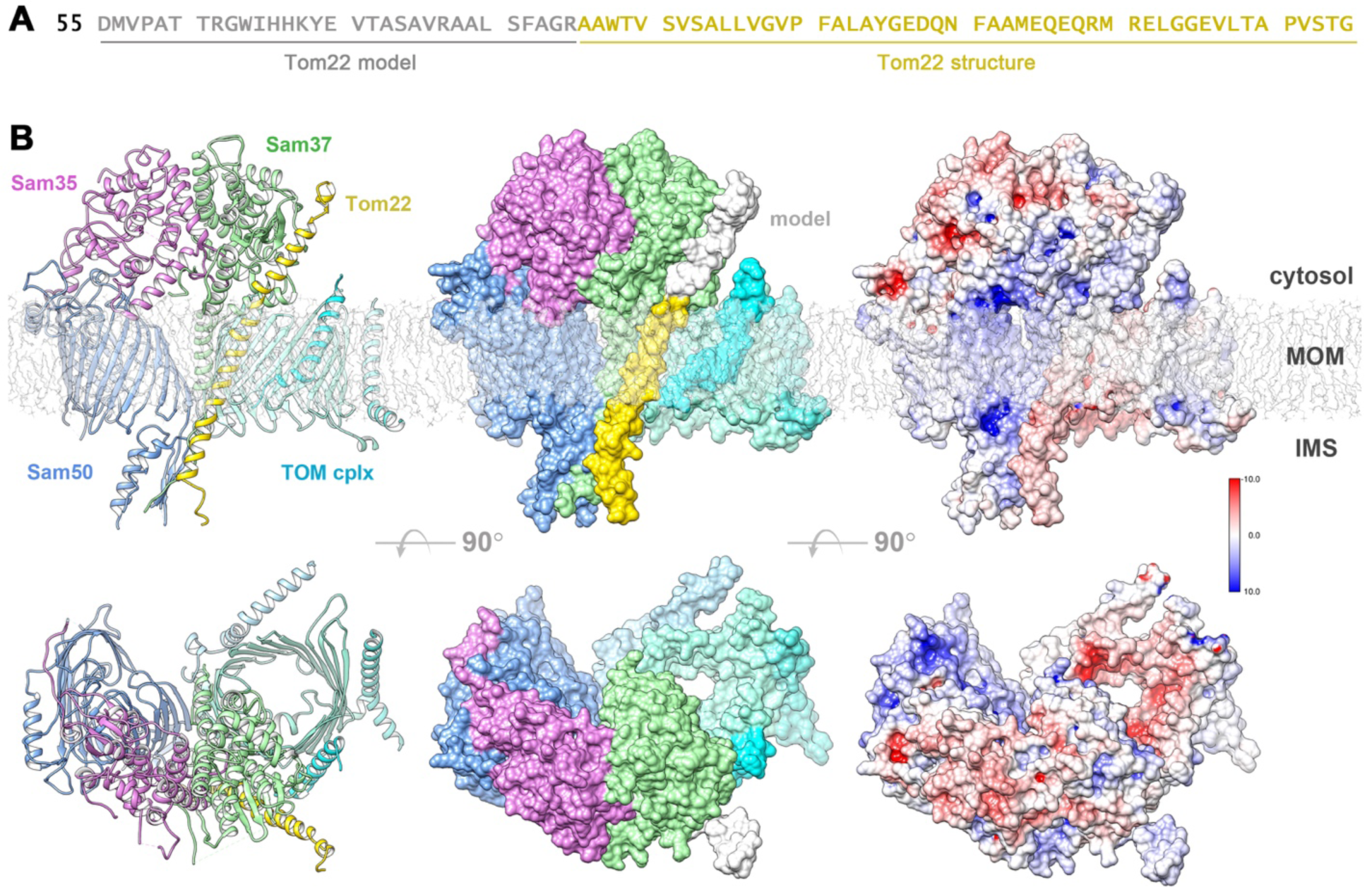
Model of the SAM-TOM supercomplex. (A) Sequence of *Mt*Tom22 used for modeling with the *ab-initio* modeled part (grey) and the homologous *Sc*Tom22 structure (gold). (B) Cartoon representation, molecular surface and surface charges for the SAM-TOM supercomplex viewed from the membrane plane and from the cytosol (MOM = Mitochondrial outer membrane, IMS = Intermembrane space). *Mt*Tom22 cartoon is shown in gold while the *Mt*Tom22 molecular surface shows the transmembrane helix (gold) homologous to the *Sc*Tom22 structure (PDB:6ucu, 6jnf) and the modeled part (grey). This Sam37-*Mt*Tom22 interaction mode perfectly accommodates the rest of the SAM and TOM complexes without clashes and the proper orientation in the mitochondrial membrane. See also Supplementary Table 6.

In the detergent structure, the Sam37 linker between transmembrane domains interacts with the Sam50 POTRA domain, but neither of these regions has high sequence conservation between species (Supplementary Fig. 6D). The majority of Sam50 L6 and β-barrel interactions are between highly conserved residues, or semi-conserved residues. Additionally, on the backside of the Sam50 β-barrel (opposite the β1-β16 interface), there are two vertical lines of conserved residues (one formed with residues on β11, β10 and β8 and the other formed by residues on β10-β7). Of these highly conserved residues, two histidine sidechains (H272 on β8, H236 on β7) face the membrane and could be important for interacting with other membrane proteins, such as Mdm10^32^. There are also three highly conserved residues on the intermembrane space loop between β8 and β9 of Sam50.

### Comparison of Sam50 and BamA: POTRA domains, lateral gate, and loop 6

Superposition of Sam50 with BamA from *Neisseria gonorrhea* (PDB:4K3B)^19^ shows that both proteins use a 16-stranded β-barrel to span the membrane, with one (Sam50) or five (BamA) POTRA domains attached at the N-terminus (Fig. 5A). The β-barrels superpose with an RMSD of 2.047 Å, with a conserved tilt angle (shear number) of the strands in the membrane. In Sam50, the POTRA domain is positioned away from the barrel lumen, potentially allowing substrate access in this conformation. In *N. gonorrhea* BamA, the POTRA domains are positioned to occlude access. Such differences have been observed in a variety of bacterial Omp85 structures. The Sam50 β-barrel is only partially closed, and PyMOL analysis does not identify any interactions at the lateral gate (Fig. 5A, Supplementary Fig 7A, B and Supplementary Table 4). In contrast, the *N. gonorrhea* BamA β-barrel is closed by 4 hydrogen bonds; two between strands β1 and β16, one between loop 1 and β16, and one between β1 and a residue after the C-terminal glycine kink in β16 (Fig. 5A, Supplementary Fig. 7A, B and Supplementary Table 4). Strands β1 and β16 form the lateral gate that opens and closes to accommodate substrate as it folds. An important feature of Omp85 proteins is that strand β16 contains a conserved glycine in the middle that facilitates kinking of that strand so that it can fold into the lumen of the barrel. The kink is required for lateral gating in BamA^33^, and this glycine is also conserved in Sam50 (Supplementary Fig. 5). In Sam50, we observe a similar kinking of strand β16 that likely facilitates lateral gate opening.

In Sam50, cytoplasmic loops L1, L2, and L3, fold in toward the barrel lumen to create the irregularly shaped barrel, whereas the BamA barrel is more elliptical (Fig. 5A). A number of other differences are observed for the outward facing loops. In BamA, loops L4 and L7 extend from the membrane surface and fold over the top of the barrel, forming a ‘capping dome’ that prevents substrate efflux. In Sam50, L4 and L7 adopt more open conformations and interact with the N-terminal region of Sam35; nonetheless these interactions also close the barrel lumen and prevent substrate efflux (Fig. 5B). Interestingly, loop 7 forms a 4-turn *α*-helix which sits parallel to the plane of the membrane. This has not been observed in BamA proteins and its function is unclear at present, although the position suggests that it may interact with the outer leaflet of the membrane.

In all Omp85 proteins, there is a conserved (V/I)RG(F/Y) motif on loop L6, where mutations to alanine abolish activity^18,34^. The conformation of the (V/I)RG(F/Y) motif is virtually identical for Sam50 and BamA, illustrating the strong structural and sequence conservation between bacteria and mitochondria (Fig. 5C). However, L6 is attached to the β-barrel of BamA very differently from Sam50.

In BamA, L6 interacts with strands β12 and β13 in the barrel lumen, using a highly conserved aspartate on β13 to form a salt bridge with an arginine on L6 that is also highly conserved (Supplementary Fig. 7C and Supplementary Table 5). In fact, this interaction is seen in all bacterial Omp85 structures solved to date. An additional salt bridge is present between a highly conserved glutamate on β12 and the highly conserved L6 arginine. In Sam50, both of these residues on β12 and β13 are asparagines, however only the semi-conserved β12 residue (N402) interacts with L6 in our structure. Additional hydrogen bonds are formed between BamA L6 backbone and β12 (asparagine), β13 (aspartate), β14 (serine), and β16 (glutamine). L6 tyrosine 662 forms two hydrogen bonds with the backbone of β14 (tyrosine) and L8 (lysine).

In contrast, Sam50 does not make any of these interactions between L6 and β13, or β14. Instead, a highly conserved L6 arginine forms a hydrogen bond with semi-conserved asparagine on β12 and a salt bridge with a highly conserved glutamate on β15 (Supplementary Fig. 7C and Supplementary Table 5). Sam50 forms a unique interaction with β11 between semi-conserved arginine 340 and L6 aspartate 373. The backbone of asparagine 373 forms a hydrogen bond with the C-terminal residue, leucine 512. This asparagine also forms a hydrogen bond with serine 510, which is located after the glycine kink in β16. These two interactions stabilize the kinked conformation of β16. The interaction between highly conserved residues glutamate 504 (β16) and phenylalanine 377 (L6) is also of interest. This interaction is present in *N. gonorrhoeae* BamA between highly conserved glutamic acid 785 (β16) and semi-conserved tyrosine 662 (L6). Another similarity between Sam50 and BamA is the backbone interaction between L6 and L8, which likely also serves to stabilize the L6 conformation.

Although the functions of L6 are not completely clear, it seems to play a role in β-barrel stability in BamA proteins as they open and close at the lateral gate^19^, while in Sam50 it plays an additional more active role, participating in substrate transfer to the lateral gate^23^. The interaction observed between β16 and L6 may help guide substrates to the lateral gate, explaining how substrates can interact with both L6 and β16 in mitochondria.

### Opening of the Sam50 lateral gate observed in SAM complex dimers

3D reconstructions for the SAM complex in GDN showed a mixture of monomers and three predominant dimer conformations (Supplementary Fig. 4). Among the dimer conformations, substantial differences were observed for the Sam50 β-barrel. In dimer 1 as previously discussed, the Sam50 β-barrel is partially closed. However, in dimers 2 and 3, the β-barrel is open, allowing for association of strands β1 in an antiparallel arrangement, extending the β-sheet across two Sam50 β-barrels (Supplementary Fig. 4A). While all three dimers are clearly non-physiological, they illustrate how strand β1 of Sam50 could associate with a precursor protein^23^. A morph of Sam50 structures from dimers 1 and 3 shows how strands β1-β4 undergo an approximate 45° rotation, opening the β-barrel between strands β1 and β16 to create space for precursor protein to bind. In this morph between our observed conformations, strands β5-β16 remain mostly fixed, as does loop L6 and the conserved (V/I)RG(F/Y) motif (Supplementary Movie 1). Additional structural changes are likely to occur when substrate is present. The large conformational change of strands β1-β4 is unlike that in BamA, where the whole β1-β4 region of the barrel moves vertically and laterally. In BamA structures, the POTRA domain can move together with the base of strand β1, blocking access to the β-barrel lumen and also reducing the barrel diameter on the periplasmic side. In Sam50, the barrel diameter by the POTRA domain stays constant while the upper 3/4 region of strands β1-β4 can flip outward. The POTRA domain is almost static compared to BamA. The closed conformation of the barrel in BamA is somewhat similar to the open conformation in Sam50 if we disregard the position of the POTRA domain. The POTRA domain in Sam50 never obstructs the lumen of the barrel as in BamA, being positioned more like in the open conformation of BamA.

### Molecular modeling of the TOM-SAM supercomplex

It was previously shown that the cytosolic domain of Tom22 interacts with Sam37 in the cytosol to create a TOM-SAM supercomplex^7^. More precisely, it was found that the 54 N-terminal residues of Tom22 do not take part in this interaction. Based on these results, we modeled the Sam37-Tom22 interaction using a truncated sequence for *Mt*Tom22 (residues 55-135). While the transmembrane helix of *Sc*Tom22 has a known structure^25,26^, *Mt*Tom22 residues 55-85 were modeled *ab-initio*. The resulting *Mt*Tom22 in the context of the known *Sc*TOM core complex^25,26^ is an extended transmembrane *α*-helix with very limited degrees of freedom in the truncated cytosolic domain, which is positively charged. Examining the surface charges in Sam37, we identified two negatively charged regions that could interact with *Mt*Tom22. The protein-protein interaction analysis program (see Methods for details) identified one of these regions as the Sam37-*Mt*Tom22 interaction surface. The resulting Sam37-Tom22 complex was used to superimpose the SAM and TOM complexes on it, yielding a SAM-TOM supercomplex (Fig. 6). The supercomplex has the proper orientation in the mitochondrial membrane and has no major clashes between the members of the SAM and TOM complexes. The member proteins of the two complexes fit into this supercomplex without any adjustment, further validating the correctness of this possible interaction.

Based on sequence homology, it is likely that the Tom22-Sam37 interaction occurs through the Sam37 N-terminal domain because this part of the sequence is better conserved, and not all Sam37 subunits from different species are predicted to have C-terminal transmembrane helices. This further validates the correctness of our model. Tom22 in the reported structures^25,26^ is missing the cytosolic part, suggesting that this portion is very flexible. The *Mt*Tom22 interaction with Sam37 identified by the program is rigid-body only and conformational changes in the N-terminal region of *Mt*Tom22 could occur to further enhance the interaction. Since the interaction is not optimized, we cannot characterize it in atomic detail. Fifty-three residues of Sam37 participate in the interaction, from segments 35-63 and 326-421. There are 10 H-bonds and 4 salt bridges with an interaction surface area of 1479.6 Å^2^ (Supplementary Table 6). From the Sam37 interacting residues, only region 38-48 is conserved (Fig. 6).

In the SAM-TOM supercomplex, the Tom40 β-barrel sits adjacent to the Sam50 β-barrel. In this arrangement, the second TOM complex from a TOM-TOM dimer is displaced by the SAM complex. Since the TOM complex has been proposed to form dimers and trimers to enhance its stability^35^, replacing a dissociated copy of TOM by another binding partner could be advantageous.

## Discussion

In mitochondria and bacteria, precursor proteins are recognized by the SAM or BAM folding machinery through a sorting signal in the final β-strand, called the β-signal in mitochondria^3^ and the C-terminal signature sequence in bacteria^36^. These signature motifs are related but not identical; for example, bacterial motifs almost always end with a C-terminal phenylalanine or tryptophan and do not tolerate extensions beyond this, while mitochondrial motifs place an additional residue after the final hydrophobic position. However, sufficient similarity between the two systems exists to allow mitochondrial VDAC expression in bacterial outer membranes^37^ and bacterial OMP expression in mitochondrial outer membranes^38^. BAM and SAM are clearly similar enough to recognize one another’s substrates. In both systems, the β-signal directs the precursor protein to the folding machinery and is thought to make the first interactions with it by binding at the lateral gate of Sam50 or BamA. Disulfide crosslinking experiments on SAM^23^ and BAM^20^ show that the β-signal of the precursor protein inserts into the lateral gate by binding to strand β1 on Sam50 or BamA, displacing the native Sam50 or BamA β-signal (strand β16) to form a stable interface between folding machinery and precursor protein. Precursor proteins also interact with strand β16 on the other side of the lateral gate, but these interactions are more flexible, suggesting that N-terminal portions of the precursor protein are added at this interface while the C-terminal precursor strand remains stably bound to β1 of Sam50 or BamA. Experiments on SAM show that Sam50 L6 is required for β-signal binding and for insertion of subsequent β-hairpins^23^. Our structures illustrate how Sam50 can open sufficiently at the lateral gate to accommodate precursor binding at strand β1, positioning L6 at the lateral gate through its interaction(s) with strand β16. Future experiments will explore interactions between Sam50 strands β1, β16, and L6 with a true β-signal peptide.

Sam35 specifically recognizes β-signal peptides independently of Sam50^3^, but from a structural standpoint it remains unclear how this interaction occurs. Our analysis of the semi-conserved residues on Sam35 identified a groove with increased conservation which could be a potential binding pocket. However, this region is on the cytosolic side of the membrane which poses a topological conundrum, because SAM complex substrates reside in the intermembrane space prior to their folding and insertion into the outer membrane. Additionally, the top of the Sam50 β-barrel is occluded by the cytoplasmic loops of Sam50 and the N-terminus of Sam35, so it is less likely that substrates will pass all the way through the barrel to interact with this semi-conserved groove on Sam35. While the N-terminal residues of Sam35 located across the top of Sam50 are not well conserved, they would be in a closer position to interact with substrate inside the Sam50 β-barrel.

Sam50 transiently associates with Mdm10 in the outer mitochondrial membrane to aid in TOM complex biogenesis^39^. In *S. cerevisiae* Mdm10, two conserved aromatic residues (Y73 and Y75) are important for association with the SAM complex, and mutation of these residues disrupts the Sam50-Mdm10 interaction as well as Tom40 and Tom22 biogenesis^32^. Homology models of Mdm10 predict Y73 and Y75 are located on strand β3 within the mitochondrial outer membrane^32,40^. Analysis of Sam50 sequence conservation mapped to our structure identifies two stretches of highly conserved residues on strands β7-β11, which are opposite of the lateral gate (Supplementary Fig. 6). These highly conserved residues on Sam50, particularly those on β7 and β8 (H236 and H272 respectively), are in a reasonable position to interact with the conserved Mdm10 residues important for Sam50-Mdm10 interaction. Experimental evaluation of the involvement of these highly conserved Sam50 residues in the interaction with Mdm10 will be required. It is not currently understood how the Sam50-Mdm10 interaction contributes to TOM complex biogenesis.

## Methods

### Cloning of Expression Vectors

*Myceliophthora thermophila (*recently renamed *Thermothelomyces thermophilus;* we will continue to use *M. thermophila* here) SAM complex subunit coding sequences were codon optimized for expression in *S. cerevisiae* and obtained from GenScript. Sam50, Sam35, and Sam37 genes were cloned into pBEVY expression vectors after the GAL1 promoter using In-Fusion Cloning (Takara Bio USA, Inc). The Sam37 construct included a N-terminal Twin-Strep-tag® (IBA GmbH) and a glycine-glycine linker. Each construct was verified by sequence analysis (Macrogen USA).

### Yeast Transformation and Growth

The three SAM complex pBEVY vectors were co-transformed into *Saccharomyces cerevisiae* strain W303.1B using the lithium acetate/single-strand carrier DNA/PEG method^41-43^. Transformed cells were plated on a selection agar (6.9g/L yeast nitrogen base without amino acids, 0.62g/L Clontech -Leu/-Trp/-Ura dropout supplement, 20g/L bacto agar, 2% D-(+)- glucose) and incubated for 72 hours at 30°C.

One colony was used to inoculate 10mL of selection media (6.9 g/L Yeast Nitrogen Base without amino acids, 0.62g/L Clontech -Leu/-Trp/-Ura dropout supplement, 2% D-(+)-glucose) and incubated overnight at 30°C and 220rpm. The following day, 500mL selection media in a 1L glass flask was inoculated with the 10mL starter culture and incubated overnight at 30°C and 220rpm. Twelve 2L glass flasks containing 1L YPG media (10 g/L yeast extract, 20 g/L peptone, 30 mL glycerol, 0.1% glucose) were each inoculated with starter culture to reach a start OD_600nm_ of 0.15 and incubated for 16-18 hours at 30°C and 220rpm. Expression was induced with 0.4% D-galactose (final concentration), and cultures were allowed to grow for 4 hours at 30°C and 220rpm. Cells were harvested and washed with cold sterile ultra-pure water before storing at - 80°C.

### Mitochondrial Isolation

Thawed cell pellet was resuspended in breaking buffer (650mM sorbitol, 100mM Tris-HCl, pH 8.0, 5mM EDTA, pH 8.0, 5mM amino hexanoic acid, 5mM benzamidine, 0.2% BSA) and stirred for 30 minutes at 4°C. Once resuspended, 4mL of 200mM PMSF (PMSF in 100% ethanol) was added. Resuspended cells were passed through a Dyno-Mill Multi Lab (WAB) bead mill (0.5-0.75µm glass beads) at a flow rate of 35mL/min and the chamber temperature was maintained below 10°C^44^. Disrupted cells were collected on ice and 4mL 200mM PMSF was added.

Cell debris was removed by two centrifugation steps, transferring the supernatant to fresh tubes after the first (3,470 x g, 30 minutes each, 4°C). Mitochondrial membrane sample was isolated by centrifugation (24,360 x g, 50 minutes, 4°C). The pellet was resuspended in wash buffer (650mM sorbitol, 100mM Tris-HCl, pH 7.5, 5mM amino hexanoic acid, 5mM benzamidine) using a Dounce homogenizer on ice. Sample was centrifuged again (24,360 x g, 50 minutes, 4°C). Pellet was resuspended by homogenization in Tris-buffered glycerol (TBG) mitochondrial storage buffer (100mM Tris-HCl, pH 8.0, 10% glycerol), and centrifuged once more (24,360 x g, 50 minutes, 4°C). Pellet was resuspended by homogenization in TBG storage buffer and separated into three aliquots. Protein concentration of the mitochondrial membrane sample was determined using Pierce BCA Protein Assay Kit (Thermo Fischer Scientific). Mitochondrial membrane aliquots were flash frozen in liquid nitrogen and stored at -80°C.

### SAM Complex Nanodisc Incorporation

One aliquot of mitochondrial membrane was thawed, and the protein concentration was adjusted to 10mg/mL with TBG mitochondrial storage buffer. Sodium chloride was added to a final concentration of 150mM, and Roche cOmplete Protease Inhibitor Cocktail tablets were added. Sample was mixed with stir bar at 4°C until the protease inhibitor tablets dissolved.

Membrane was solubilized by addition of 2% final concentration of LMNG (Anatrace) and stirring for 1.5 hours at 4°C. Solubilized material was isolated by ultracentrifugation (208,000 x g, 45 minutes, 4°C). The supernatant was filtered with a 0.22µm SteriFlip vacuum filter (Millipore) and used immediately.

Filtered supernatant was added to Strep-Tactin Sepharose (IBA GmbH) resin and rocked for 4 hours at 4°C. Following incubation, sample was transferred to a gravity flow column and washed with four column volumes of wash buffer (100mM Tris-HCl, pH 8.0, 150mM NaCl, 1mM EDTA, pH 8.0, 0.02% LMNG). Protein was eluted in 0.5 column volume fractions using elution buffer (100mM Tris-HCl, pH 8.0, 150mM NaCl, 1mM EDTA, pH 8.0, 2.5mM Desthiobiotin, 0.02% LMNG). Fractions were analyzed by SDS PAGE, and those containing SAM complex were pooled and concentrated using 100kDa molecular weight cut off Amicon Ultra Centrifugal Filter Unit (Millipore).

Concentrated sample was injected onto HiLoad 16/600 Superose 6 prep grade column (GE Healthcare) at a flow rate of 0.11mL/min, in size-exclusion buffer (20mM HEPES, pH 8.0, 150mM NaCl, 0.02% LMNG). Fractions were collected and analyzed by SDS-PAGE.

MSP1E3D1 plasmid (Addgene) was expressed and purified as previously described^45^. Purified SAM complex in LMNG, MSP1E3D1, and cholate solubilized DOPE (Avanti Polar Lipids, Inc.) and DOPC (Anatrace) were mixed in a molar ratio of 1 SAM : 2 MSP1E3D1 : 200 DOPE : 200 DOPC. Additional cholate was added to obtain final concentration of 20mM. Mixture was incubated for one hour on ice, then 0.1g Bio-Beads SM2 (BioRad) were added per milliliter of incorporation sample to facilitate nanodisc formation. Sample with Bio-Beads was rocked at 4°C overnight. An additional 0.1g/mL Bio-Beads SM2 were added the following morning and sample was rocked for one hour at 4°C. Bio-Beads were removed from sample with low speed centrifugation. Insoluble protein was separated by ultracentrifugation (208,000 x g, 45 minutes, 4°C).

Soluble fraction was added to Strep-Tactin Sepharose (IBA GmbH) resin and rocked for 4 hours at 4°C. Sample was transferred to a gravity flow column and washed with wash buffer (100mM Tris-HCl, pH 8.0, 150mM NaCl, 1mM EDTA, pH 8.0). SAM complex in nanodiscs was eluted with elution buffer (100mM Tris-HCl, pH 8.0, 150mM NaCl, 1mM EDTA, pH 8.0, 2.5mM Desthiobiotin). Fractions were analyzed with SDS-PAGE and those containing SAM complex in nanodiscs were pooled and concentrated to 1mL using 100kDa molecular weight cut off Amicon Ultra Centrifugal Filter Unit (Millipore).

Sample was injected onto Superose 6 10/300 column (GE Healthcare) at a flow rate of 0.20mL/min in size-exclusion buffer (20mM HEPES, pH 8.0, 150mM NaCl). Fractions were collected and analyzed by SDS-PAGE. The peak fraction containing SAM complex in nanodisc was concentrated to ∼1mg/mL for cryo-electron microscopy.

### SAM Complex Purification in Detergent

SAM complex was purified as described above, but detergent was exchanged to 0.02% GDN (Anatrace) during size-exclusion chromatography. Briefly, isolated mitochondria were solubilized in 2% LMNG for 1.5 hours at 4°C. Filtered soluble fraction was bound to Strep-Tactin Sepharose (IBA GmbH) resin for 4 hours, then SAM complex was eluted with Desthiobiotin elution buffer. Concentrated sample was injected onto HiLoad 16/600 Superose 6 prep grade column (GE Healthcare) at a flow rate of 0.11mL/min, in size-exclusion buffer (20mM HEPES, pH 8.0, 150mM NaCl, 0.02% GDN). Fractions were collected and analyzed by SDS-PAGE. Peak fractions containing SAM complex were pooled and concentrated to ∼5mg/mL for cryo-electron microscopy.

### CryoEM sample preparation and data collection

An aliquot of 3 μL of SAM complex sample was applied to freshly glow-discharged holey carbon grid (Quantifoil R1.2/1.3, copper, 300 mesh). The grids were blotted for 6 s and plunge-frozen in liquid nitrogen-cooled liquid ethane using an FEI Vitrobot Mark IV plunger. CryoEM data were collected on a Titan Krios G3 electron microscope (Thermo-Fisher) operated at 300kV and equipped with a Gatan Quantum LS imaging energy filter with the slit width set at 20 eV. Micrographs were acquired on a K2 Summit direct electron detection camera at the nominal magnification of 130,000x (calibrated pixel size of 1.06 Å on the sample level) using the Leginon automation software package ^46^. The dose rate on the camera was set to 8 e^−^/pixel/s. The total exposure time of each micrograph was 10 s fractionated into 50 frames with 0.2 s exposure time for each frame. Detailed data collection parameters are listed in Table 1.

### Image processing

All frames of each dose-fractionated micrograph were aligned for correction of beam-induced drift using MotionCor2^47^. Two average images were generated from motion correction for each micrograph: one with dose weighting and the other one without. The average images without dose weighting were used for defocus determination using CTFFIND4^48^. Quality of the micrographs was evaluated using the results from CTFFIND4. The micrographs with poor resolution (worse than 4.5 Å) or too large (>3.0 µm) or too small (<0.8 µm) underfocus values were removed. The single particle data analysis was performed following the standard procedures in RELION3^49^ and cryoSPARC2^50^ with few modifications as summarized in the following and in Supplementary Figure 1C and 2.

### SAM complex in lipid nanodiscs

The procedures are summarized in Figure S1C. No symmetry was applied in the data processing of the SAM complex in lipid nanodiscs. A total of 14,032 micrographs were acquired and 11,347 micrographs were selected for data processing using the results from CTFFIND4. Initially, a set of 500 micrographs were used for particle picking by Gautomatch (https://www.mrc-lmb.cam.ac.uk/kzhang/Gautomatch/). No external templates were supplied and Gautomatch generated templates automatically. The particles were processed using cryoSPARC2 for 2D classification and ab-initio reconstruction to generate starting models. The projections of the best starting model was then used as templates for particle picking by Gautomatch using all micrographs. A total of 3,951,406 particles were picked and extracted in 96×96 pixels with 2x binning (pixel size 2.12 Å). The particles were processed using RELION3 for 2D and 3D classifications. The best particles were selected iteratively by selecting the 2D class averages and 3D reconstructions that had interpretable structural features. 446,747 particles were selected using the above procedures and subjected to 3D classification with only 1 class to align particles to the center. The result was used to re-center the particles as well as to remove overlapping particles (deduplication), leading to 445,569 particles being re-extracted in 192×192 pixels without binning (pixel size 1.06 Å). After further 2D and 3D classifications, 205,131 particles were selected for 3D autorefinement to obtain a reconstruction at 4.2 Å resolution. The result was then used for particle polishing that included all frames. A 2D classification was always carried out after each run of particle polishing to remove particles containing bad pixels from the camera in the processing of all the data sets reported here. The resolution of 3D autorefinement was improved to 4.0 Å with 190,078 particles after particle polishing. Ctf refinement was then performed to refine per-particle defocus values but no resolution improvement was noticed at this step. An additional run of particles polishing was carried out using the first 30 frames and 179,509 “shiny” particles were generated for the final 3D autorefinement using RELION3 or cryoSPARC2. The refinement using RELION3 reported 3.9 Å resolution and the non-uniform refinement using cryoSPARC2 reported 3.4 Å resolution. A soft mask was used in the RELION3 3D autorefinements. The cryoSPARC2 non-uniform refinement also employed an auto-generated soft mask. In our case, the cryoSPARC2 non-uniform refinement generated better results as reported by FSC and the quality of the density maps, therefore the 3D reconstructions from cryoSPARC2 were used for structural interpretation and atomic modeling.

### SAM complex in detergent

The procedures are summarized in Figure S2. A total of 14,329 micrographs were acquired and 10,831 micrographs were selected using the results from CTFFIND4. Similar to the above procedures, a small subset of micrographs were used for particle picking without a template. The particles were processed for 2D classification using RELION3. Representative 2D class averages were selected to serve as templates for a new iteration of particle picking by Gautomatch using all the micrographs. A total of 5,842,605 particles were picked and extracted in 104×104 pixels with 2x binning (pixel size 2.12 Å). The particles were processed for 2D classifications using RELION3. The best particles were selected iteratively from the results of 2D classification. Comparison of the 2D class averages suggested that there are two different forms of complexes in the sample: “monomeric” and “dimeric” complexes (Fig. S2). These two different forms of particles were separated (514,800 “monomer” particles and 829,206 “dimer” particles) for the following processing procedures.

The processing of “monomer” particles is summarized in Figure S2. After re-centering and deduplication, 511,249 “monomer” particles were re-extracted in 208×208 pixels without binning (pixel size 1.06 Å) and subjected to 2D classification using RELION3, from which 457,279 particles were selected for further processing using cryoSPARC2. No symmetry was applied in the data processing of the “monomer” particles. After particle selection from 2D classification and 3D hetero refinement, 138,575 particles were used in non-uniform refinement to achieve a final reconstruction at 3.7 Å resolution.

The processing of “dimer” particles is summarized in Figure S2. After re-centering and deduplication, 821,025 “dimer” particles were re-extracted in 208×208 pixels without binning (pixel size 1.06 Å) and subjected to 2D and 3D classifications using RELION3. A starting model of the “dimer” was generated using the ab-initio reconstruction in cryoSPARC2 to serve as the reference map for the 3D classification in RELION3. The C2 symmetry was applied in all 3D reconstructions during the data processing of the “dimer” particles. A total of 5 classes of 3D reconstructions were generated by 3D classification. 3 of the 5 classes show interpretable structural details and their conformations are notably different, therefore the particles in these 3 classes were separately sent to cryoSPARC2 for further hetero refinement and non-uniform refinement to obtain final 3D reconstructions. It is worth noting that the particles of “dimer 1” were processed using RELION3 for 3D autorefinement, Ctf refinement, and particle polishing before the cryoSPARC2 processing. The resolutions of the 3D reconstructions of these dimers are 3.2 Å (“dimer 1” with 117,339 particles), 3.6 Å (“dimer 2” with 60,472 particles), and 3.9 Å (“dimer 3” with 122,361 particles), respectively.

The interfaces between two “monomers” in the “dimers” are relatively small and suspectable to cause flexibilities in the “dimers” as evidenced by poor densities in some regions of the 3D reconstructions (Fig. S3D). Therefore, the reconstruction of “dimer 1”, which is at the highest resolution among three different “dimers”, was further improved by dividing the “dimer” into two independent “monomers” for refinement using symmetry expansion (Fig. S3D). Briefly, 174,217 polished particles of “dimer 1” were processed using RELION3 for Ctf refinement and 3D autorefinement with the C2 symmetry. Each particle was then expanded into two particles using the program *relion_particle_symmetry_expand* to generate a total of 335,670 particles. No symmetry was applied in the afterward procedures. A soft mask, that included one “monomer” and the detergent micelle, was carefully constructed following the instructions ^51^ for density subtraction. Another soft mask that only included the remaining “monomer” after density subtraction was also constructed for the following 3D refinements. During RELION3 3D autorefinement, local search was forced by setting both parameters “initial angular sampling” and “local searches from auto-sampling” to 1.8°. A 3D reconstruction of the “monomer” was obtained at 3.3 Å resolution after the first run of 3D autorefinement using the density-subtracted particles. Then the particles were subjected to particle polishing for a second time using the first 30 frames. It is worth noting that the particles after this step of particle polishing were intact “dimers” without density subtraction. The soft mask and local search were used in the 3D refinement of the “monomer” with the polished particles of “dimer” to ensure the cross interferences between two “monomers” within each particle was minimal. Finally, the polished “shiny” particles were sent to cryoSPARC2 for a final 3D refinement with the features of local refinement and non-uniform refinement turned on to achieve the 3D reconstruction of the monomer at 3.0 Å resolution. The resolution and quality of the reconstruction were both improved by symmetry expansion as shown in Figure S3D.

### Model building and refinement

A model was manually built into the final density map of the SAM complex in lipid nanodiscs using COOT^52^. The protein sequences of *M. thermophila* Sam35, Sam37 and Sam50 (Uniprot: G2QAT9, G2Q6R7, and G2QFF9 respectively) were used to build models from scratch.

Secondary structure predictions were performed using I-TASSER^53^ and Phyre2^54^. At the level of 3.4 Å resolution, the density was sufficient to allow *ab initio* tracing of a majority of the folds of all three subunits. This model was later used as an initial reference for model building into the density maps of dimeric SAM complexes in detergent. In the highest resolution (3 Å) map of the monomeric SAM complex derived from dimer 1, a majority of the protein sidechains were well resolved, allowing model building without ambiguity. All the models were refined using both Rosetta^55^ and the real-space refinement in PHENIX^56^. The statistics are summarized in Table 1.

### Sequence Alignment

Sequences for Sam50 from different species were obtained from UniProtKB^57^. Multiple sequence alignment of Sam50 was completed using T-Coffee Expresso^58-62^. Single residue manual adjustments to the sequence alignment were completed in Jalview^63^. The final alignment with sequence similarities highlighted was produced with ESPript 3.0, with sequence similarities depiction parameters set to %Equivalent, global score of 0.6, and flashy color scheme output^64^. Lastly, the high-resolution structure from SAM complex in detergent was used to assign secondary structure to the alignment.

### Modeling the Sam37-Tom22 interaction

Since the 54 N-terminal residues of Tom22 have been shown not to be involved in Sam37 binding^7^, the *Myceliophthora thermophila* Tom22 (*Mt*Tom22; Uniprot ID: G2QBG3) sequence was truncated to 55-135 for modeling. Residues 55-85 had to be modeled *ab-initio*, while the 86-135 segment was homology modeled from the *Saccharomyces* Tom22 structure extracted from the two available TOM core complex structures (PDB:6ucu, 6jnf)^25,26^. The *Mt*Tom22 model was built by DMPfold^65^ on the PSIPRED server^66^. The resulting model was used for protein-protein interaction prediction with the standalone version of PIPER^67^. *Mt*Tom22-Sam37 complexes were visually inspected to select the *Mt*Tom22 conformations that have the correct orientation in the mitochondrial membrane, do not clash with the other members of the SAM complex and can also accommodate the other members of the TOM complex without clashing with the SAM complex. The best result was used for calculating surface charges in Chimera, using the AMBER forcefield and APBS^68^.

### Interaction analyses

The interaction analyses for Sam50/Sam35, Sam35/Sam37, and Sam50/37, as well as for Sam37-Tom22, were completed using QT Pisa^69^ and PyMOL (Version 2.3 Schrödinger, LLC), and figures were made using Chimera^70,71^.

## Data Availability

Atomic coordinates and structure factors for SAM complex structures have been deposited in the EMDB and wwPDB under accession codes EMD XXXX and PDB YYYY. Source data for all figures and files is available from the authors upon request.

## Acknowledgements

This work utilized the NIH Multi-Institute Cryo-EM Facility (MICEF) and the computational resources of the NIH HPC Biowulf cluster (http://hpc.nih.gov). We thank Huaibin Wang and Haifeng He for technical support on the NIH MICEF Titan Krios Electron Microscope. K.A.D, S.E.R., I.B, and S.K.B. are supported by the Intramural Research Program of the NIH, NIDDK. X.N., X.T., and J.J. are supported by the Intramural Research Program of the NIH, NHLBI. S.R. was also supported by a Sir Henry Wellcome Postdoctoral Fellowship. M.S.K. and E.R.S.K. are funded by program grant MC_UU_00015/1 of the Medical Research Council.

## Author contributions

K.A.D, S.E.R., and S.K.B. designed the study. K.A.D and S.E.R. cloned, expressed and purified SAM complexes. X.N. and J.J. collected and processed cryoEM data. X.N. and X.T. built the initial models. X.N, I.B. and K.A.D. refined the final models. All authors analyzed the data and wrote the manuscript.

## Declaration of interests

Authors declare no competing interests.

## Notes

### Competing Interest Statement

The authors have declared no competing interest.

