## supplemental figures for "Structural Insight into Mitochondrial β-Barrel Outer Membrane Protein Biogenesis"

### Supplementary Information

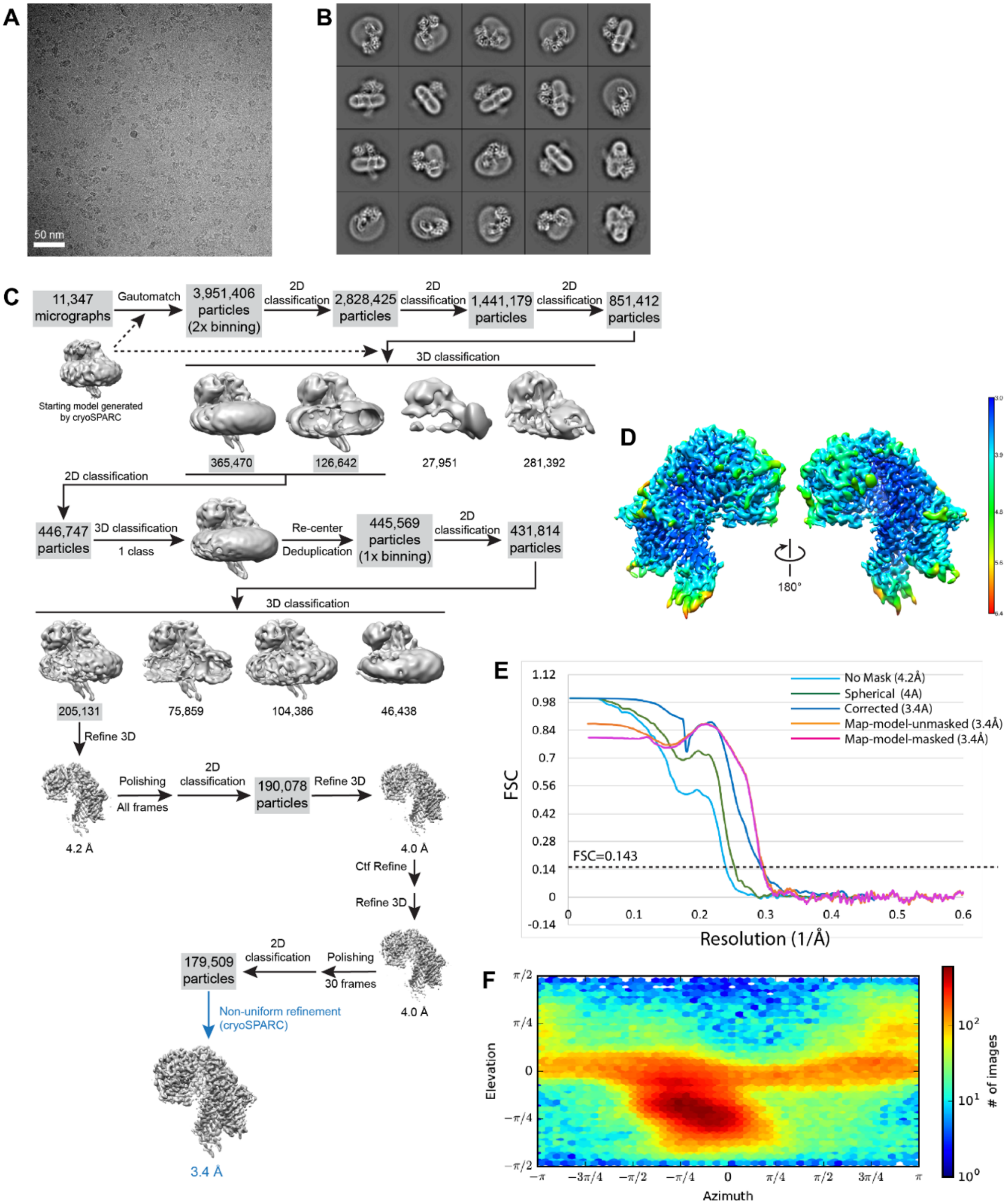

**Supplementary Figure 1. CryoEM data processing of the SAM complex in lipid nanodiscs. A.** CryoEM micrograph of the SAM complex in lipid nanodiscs. **B.** Representative 2D class averages of the SAM complex. **C.** Schematic diagram of cryoEM data processing procedures for the SAM complex in lipid nanodiscs. **D.** Local resolution map calculated by cryoSPARC2**. E.** Fourier Shell Coefficient (FSC) curves. **F.** Orientation distribution plot.

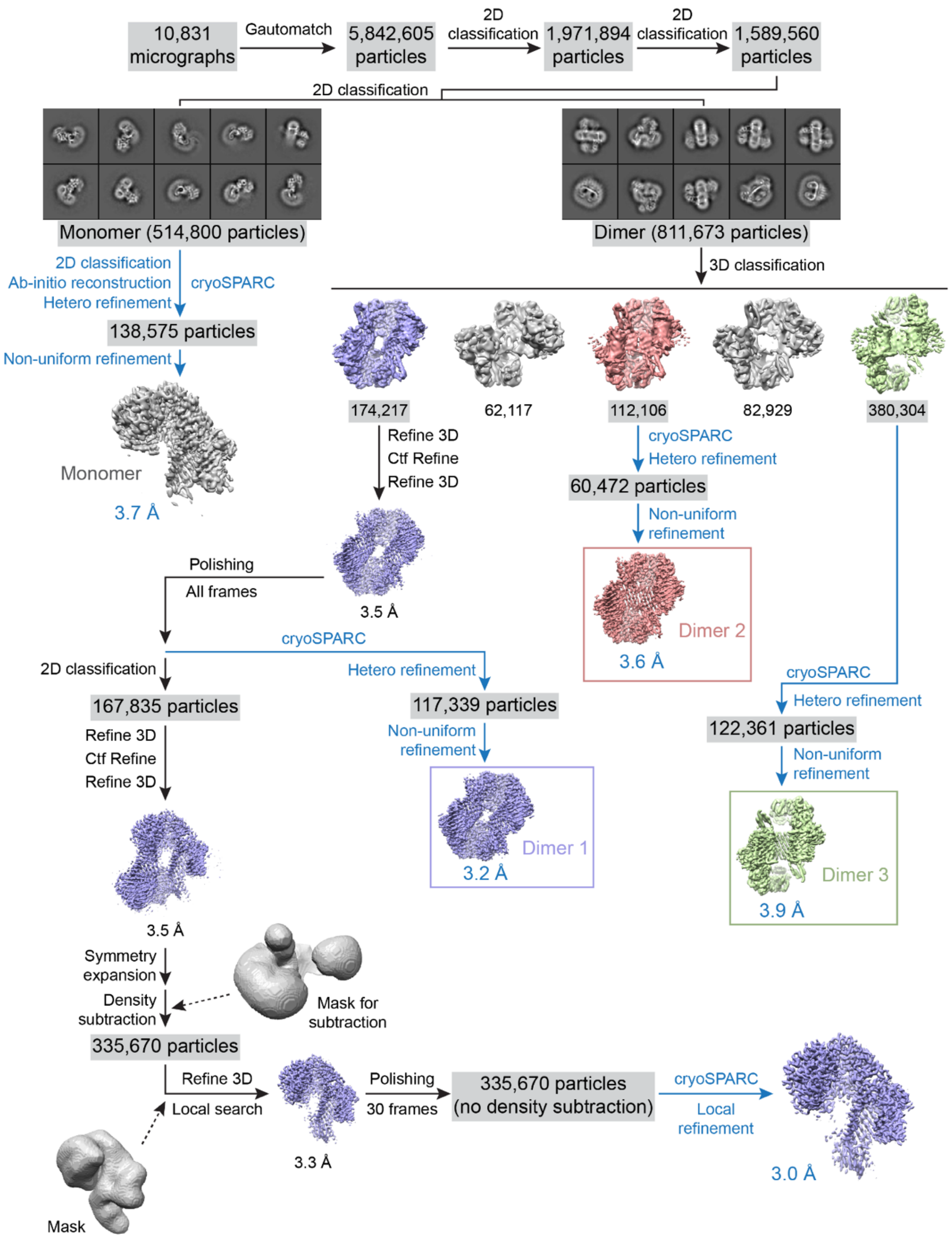

**Supplementary Figure 2. CryoEM data processing of the SAM complex in detergent GDN.** The procedures carried out using different software packages are depicted in black for RELION3 and blue for cryoSPARC2. The particles of "monomer" and "dimer" were sorted by 2D classification.

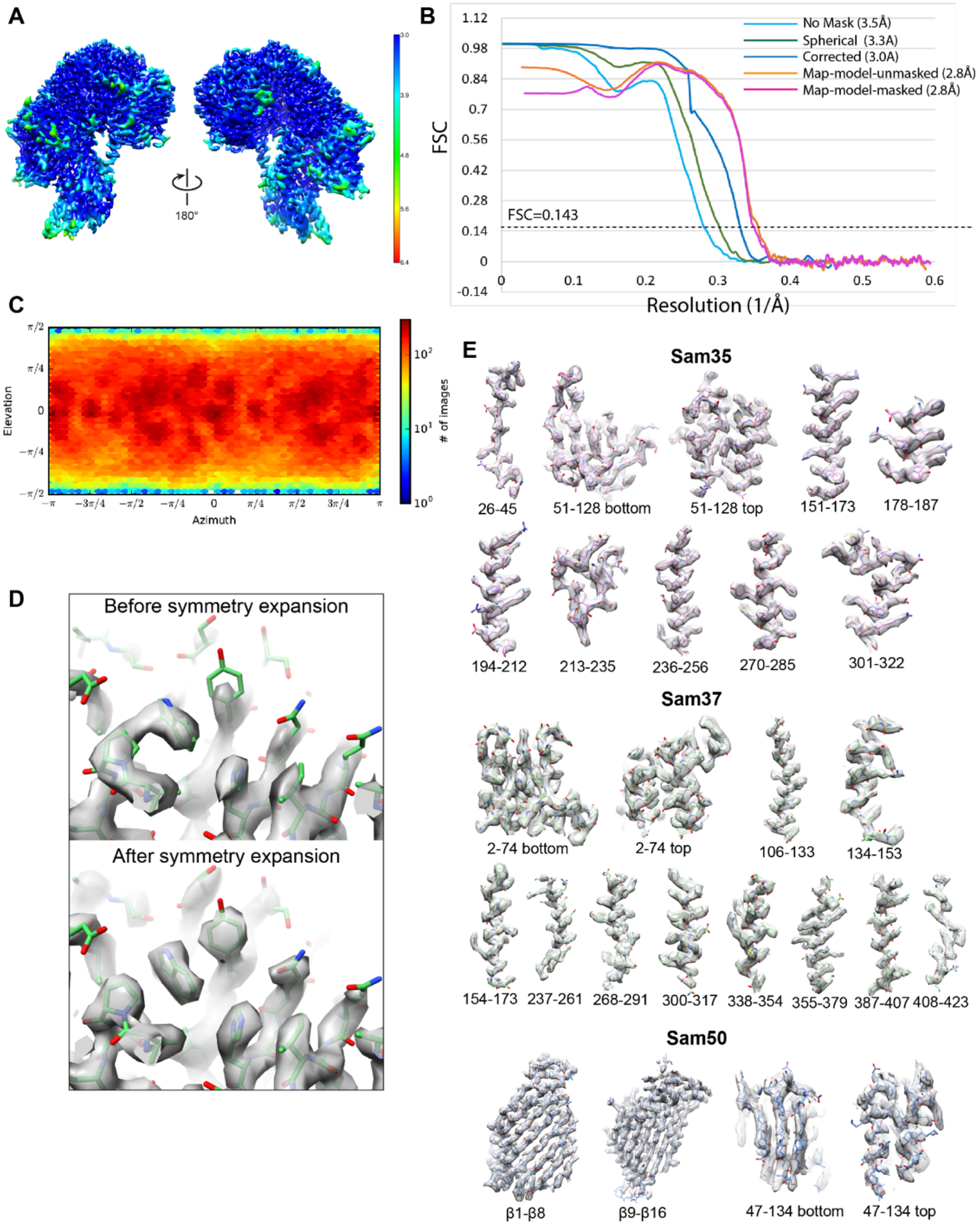

**Supplementary Figure 3. Resolution estimation of cryoEM single particle reconstruction of the SAM complex in detergent GDN with symmetry expansion. A.** Local resolution map calculated by cryoSPARC2**. B.** FSC curves. **C.** Orientation distribution plot. **D.** The resolution and quality of the cryoEM density map were both improved after symmetry expansion as demonstrated in a representative region in Sam37. **E.** Superposition of the cryoEM densities and atomic model of the SAM complex in detergent GDN.

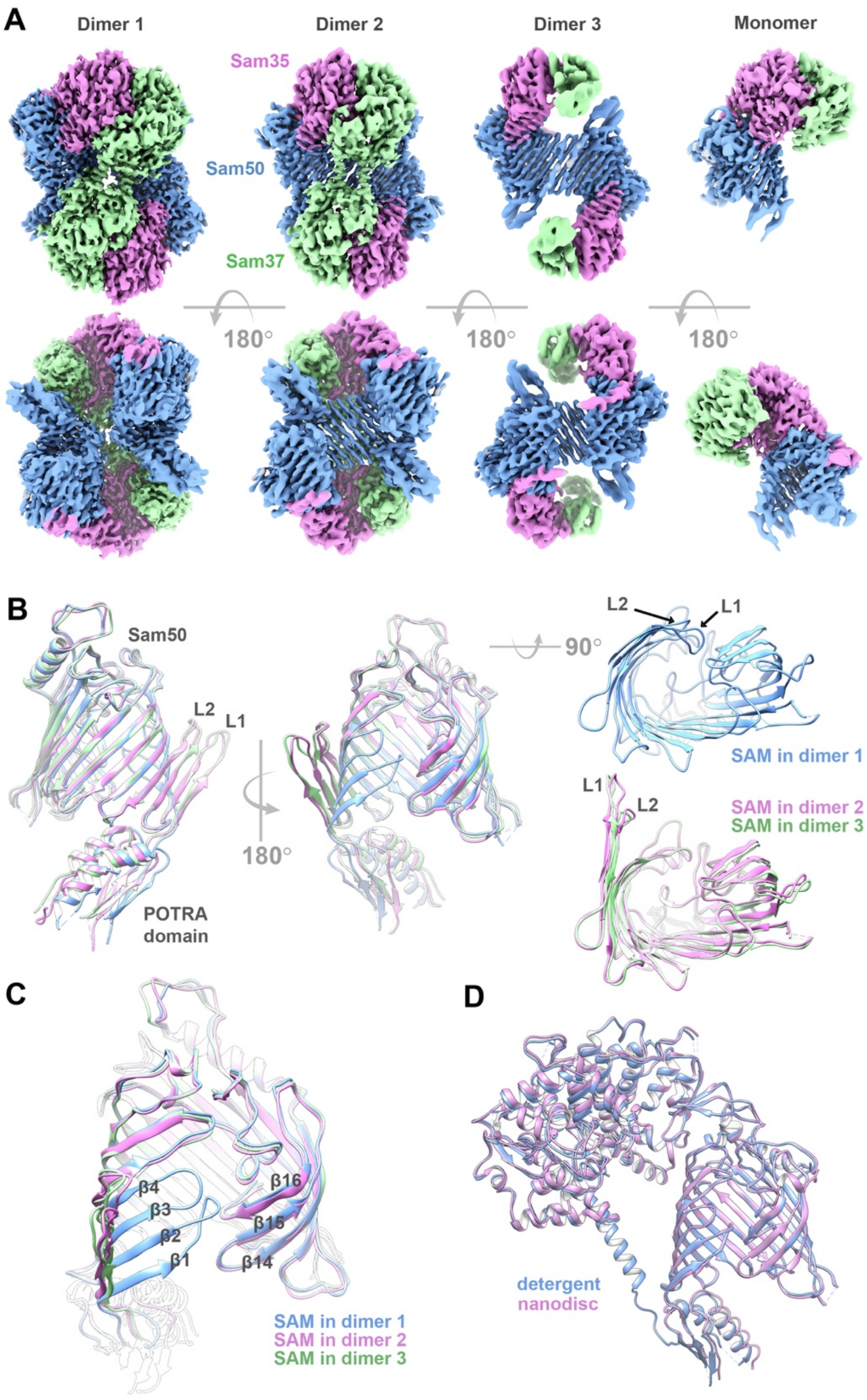

**Supplementary Figure 4. Conformations of Sam50 in detergent GDN and in lipid nanodiscs. A.** Comparison of the cryoEM density maps of the SAM complex as dimers and monomer in GDN. **B.** Superposition of Sam50 from dimer 1 (blue), dimer 2 (orchid), and dimer 3 (green). Dimer 2 and dimer 3 exhibit a barrel opened by β1-β4. **C.** Comparison of the lateral gate of Sam50 in each dimer. **D.** Comparison of the similar structures of the SAM complex in GDN (blue) and lipid nanodiscs (orchid).

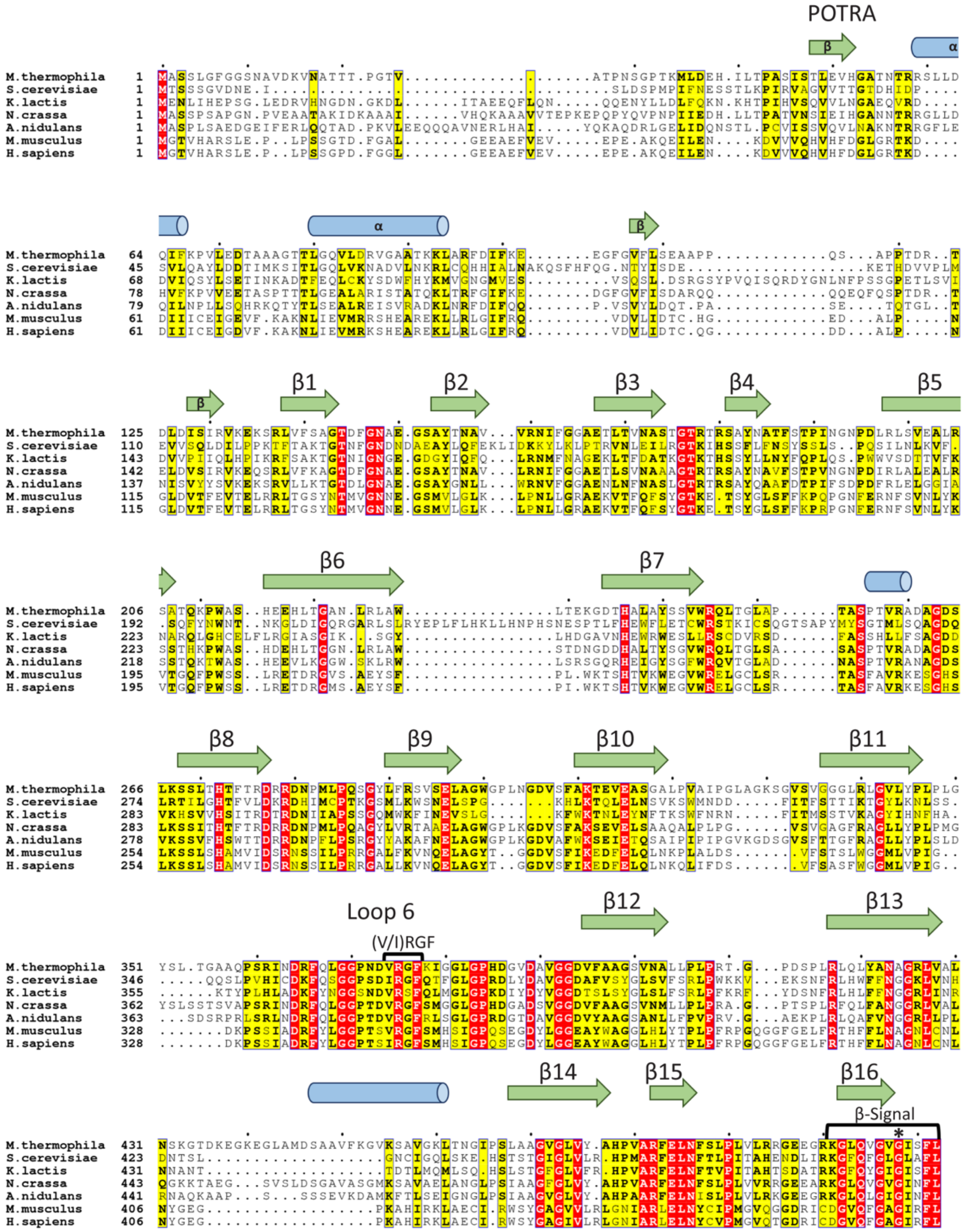

**Supplementary Figure 5. Structure-coded sequence alignment for Sam50.** Secondary structure assigned based on high-resolution structure of *M. thermophila* SAM complex in detergent. Blue cylinders represent α-helices, green arrows represent β-strands. Asterisk identifies glycine that forms a kink in β16. *M.thermophila* (*Myceliophthora thermophila,* Uniprot: G2QFF9), *S.cerevisiae* (*Saccharomyces cerevisiae,* Uniprot: P53969), *K.lactis* (*Kluyveromyces lactis,* Uniprot: Q6CNZ6), *N.crassa* (*Neurospora crassa,* Uniprot: V5IKW7), *A.nidulans* (*Aspergillus nidulans,* Uniprot: C8VCB1), *M.musculus* (*Mus musculus,* Uniprot: Q8BGH2), *H.sapiens* (*Homo sapiens,* Uniprot: Q9Y512)

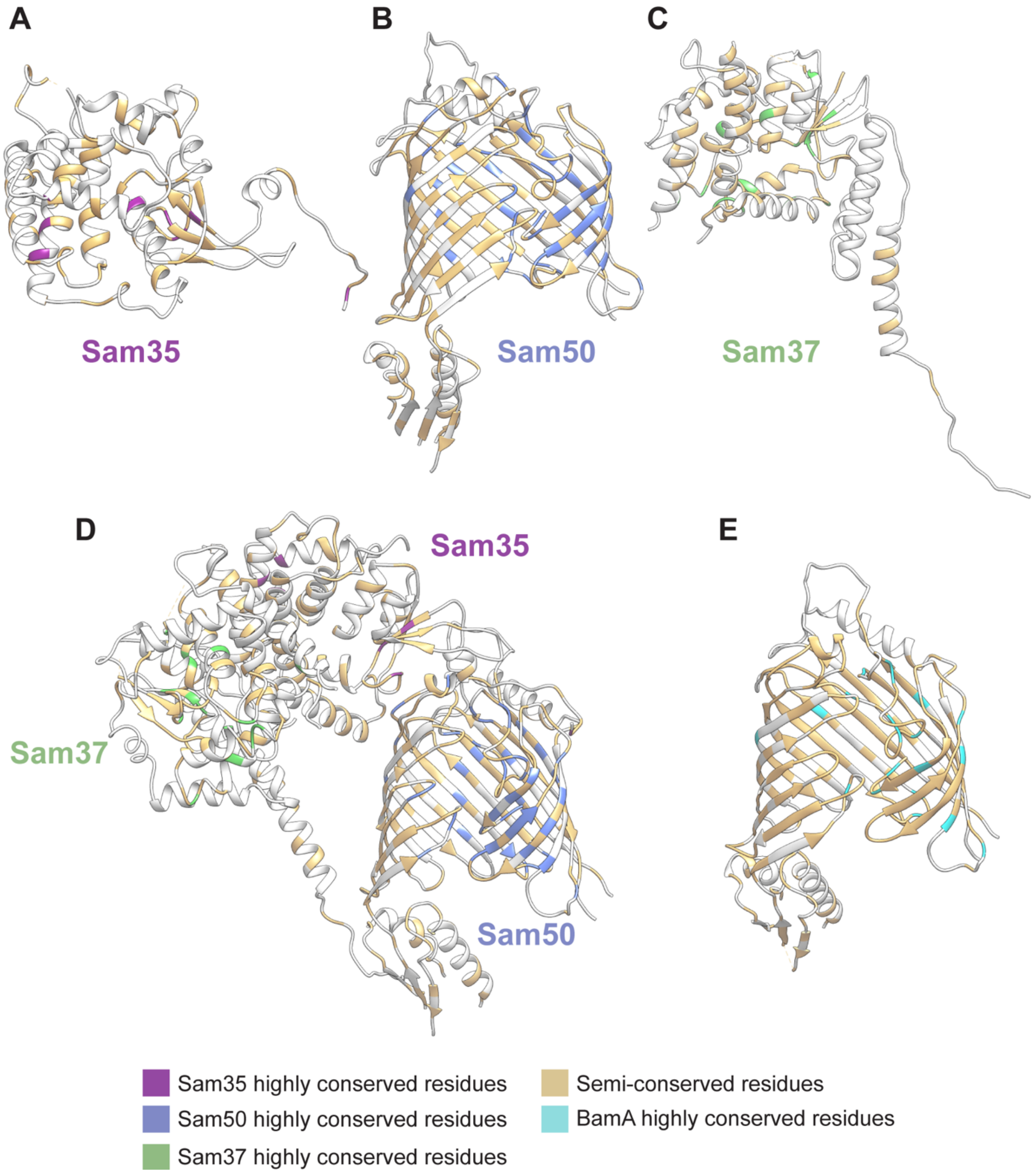

**Supplementary Figure 6. Highly conserved residues in each subunit of the SAM complex. A.** Sam35. **B.** Sam50. **C.** Sam37. **D.** The entire SAM complex. **E.** Sam50 residue conservation with *Neisseria gonorrhoeae* BamA. Highly conserved residues are in colors specific for each subunit and semi-conserved residues are in gold for all subunits.

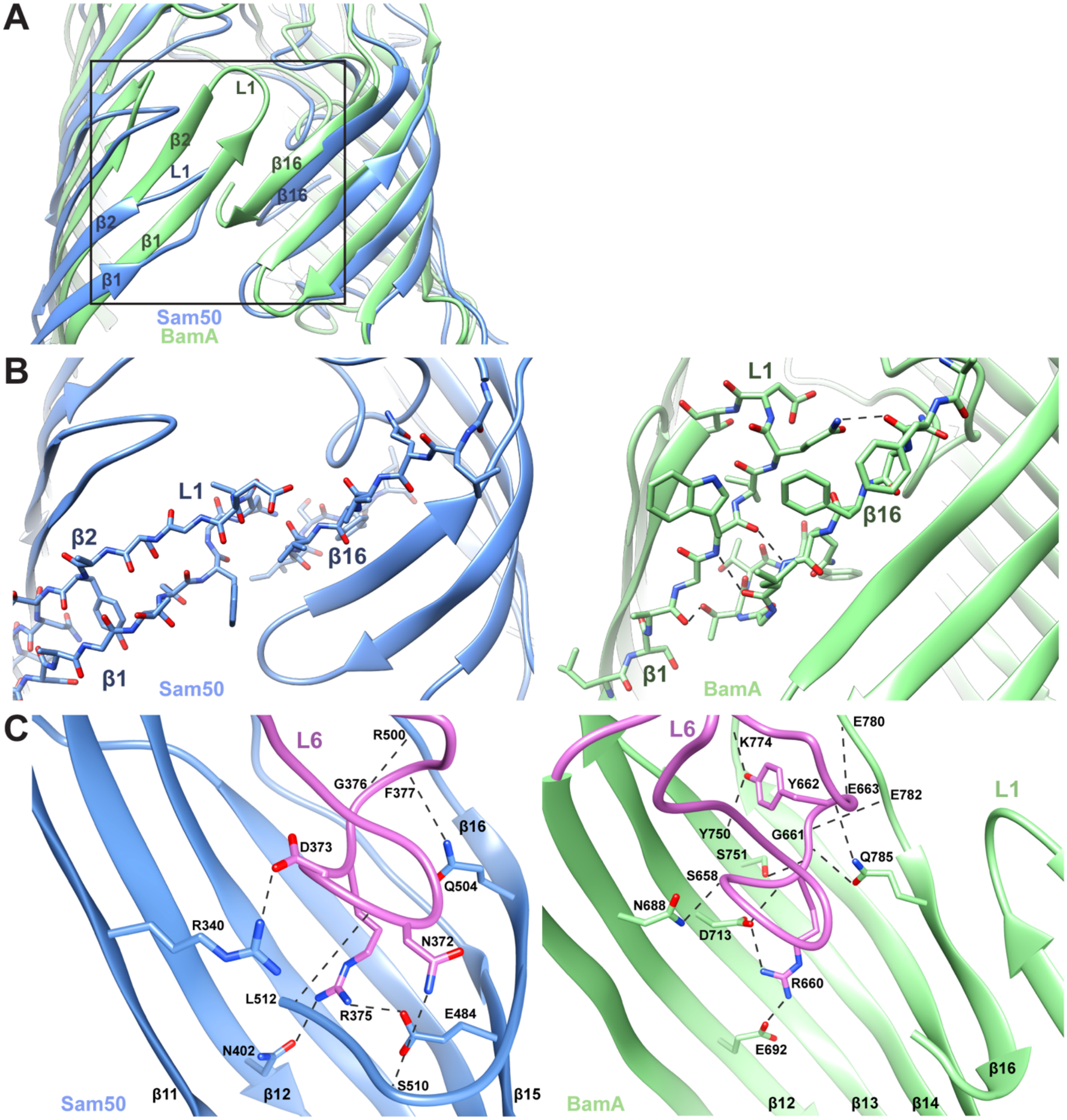

**Supplementary Figure 7. Interactions between β1 and 16 in Sam50 and BamA, and a comparison of loop 6 interactions. A.** Comparison of the lateral gate in Sam50 (blue) and BamA (green)**.** Strands β1-β2, β16 and loop 1 are indicated. The boxed area is zoomed in for each protein in [B]. **B.** Lateral gate of Sam50 on the left, and *Neisseria* BamA (PDB:4K3B) on the right. Hydrogen bonds are shown as dashed lines. **C.** Comparison of loop 6 interactions in Sam50 on the left, and *Neisseria* BamA (PDB:4K3B) on the right. Loop 6 is colored orchid in both structures, hydrogen bonds are shown as dashed lines.

| Protein | Mt Sam35 | | Mt Sam50 | | Mt Sam37 | |
|---|---|---|---|---|---|---|
| | Sam35 | Sam50 | Sam50 | Sam37 | Sam37 | Sam35 |
| **Nr of atoms** | | | | | | |
| in interface | 183 (8.0%) | 219 (6.6%) | 113 (3.4%) | 98 (3.7%) | 213 (8.1%) | 209 (9.1%) |
| on the surface | 1433 (62.4%) | | 2089 (63.4%) | | 1632 (62.1%) | |
| Total | 2295 (100.0%) | | 3295 (100.0%) | | 2628 (100.0%) | |
| **Buried Surface area ($A^2$)** | 1,972.8 (12.0%) | 1,865.4 (8.0%) | 924.4 (4.0%) | 966.9 (5.0%) | 2,011.6 (10.4%) | 1945.8 (11.8%) |
| **Accessible Surface area ($A^2$)** | 16,466.6 | | 23,272.2 | | 19,312.1 | |
| **Solvation Energy (kcal/mol)** | -19.9 | | -7.9 | | -14.3 | |
| Salt bridges in interface | 3 | | 0 | | 2 | |
| H-bonds in interface | 15 | | 8 | | 9 | |
| **Volume ($A^3$)** | 36,180 | | 52,820 | | 41,670 | |
| **Buried Surface area in cplx ($A^2$)** | 3,918.6 | | 2,789.8 | | 2,978.5 | |
| **Total surface area of cplx ($A^2$)** | 49,920 | | | | | |
| **Total buried area in cplx ($A^2$)** | 9,686.9 | | | | | |

**Supplementary Table 1. QT-Pisa analysis of molecular interfaces in the high-resolution detergent SAM complex.** Volume value from Chimera analysis, number of salt bridges and hydrogen bonds in interface from PyMOL analysis.

| Hydrogen bonds | | | | Salt bridges | | |
|---|---|---|---|---|---|---|
| **Sam37** | **distance** | **Sam35** | | **Sam37** | **distance** | **Sam35** |
| H-28 N | 3.3 | L174 O | | D111 OD1 | 3.3 | R152 NE |
| A-31 O | 3.1 | A231 N | | R363 NH2 | 2.4 | D229 OD2 |
| T49 O | 2.4 | K221 NZ | | | | |
| Q50 O | 3.1 | I222 N | | | | |
| S69 OG | 3.4 | R241 NH2 | | | | |
| D111 OD1 | 3.1 | Y156 OH | | | | |
| R168 NH1 | 3.4 | L225 O | | | | |
| E353 OE2 | 3.5 | L225 N | | | | |
| K370 NZ | 2.4 | I234 O | | | | |
| **Sam37** | **distance** | **Sam50** | | **Sam37** | **distance** | **Sam50** |
| F406 O | 2.5 | R137 NH2 | | | | |
| F413 N | 3.2 | T55 O | | | | |
| G414 N | 2.6 | T55 O | | | | |
| A416 O | 3.4 | G53 N | | | | |
| Q418 O | 3.5 | V51 N | | | | |
| Q418 NE2 | 3.3 | A54 O | | | | |
| V419 O | 3.0 | R59 NH1 | | | | |
| V419 O | 3.4 | R59 NH2 | | | | |
| **Sam50** | **distance** | **Sam35** | | **Sam50** | **distance** | **Sam35** |
| R259 NH1 | 2.6 | S95 O | | D386 OD1 | 2.9 | R108 NH1 |
| A260 O | 3.3 | R40 NH2 | | D386 OD2 | 2.8 | R108 NH2 |
| A262 O | 2.7 | T92 OG1 | | K433 NZ | 3.4 | E36 OE2 |
| *G383 O* | 3.3 | N35 ND2 | | | | |
| G387 N | 3.0 | T112 O | | | | |
| L430 O | 3.1 | E39 N | | | | |
| N431 ND2 | 2.7 | E36 OE1 | | | | |
| S432 N | 2.7 | E39 OE1 | | | | |
| S432 OG | 2.9 | E39 OE2 | | | | |
| L491 O | 2.7 | L29 N | | | | |
| L491 O | 3.5 | R30 N | | | | |
| L493 O | 3.2 | Y32 N | | | | |
| L493 N | 2.8 | R30 O | | | | |
| R495 N | 3.1 | Y32 O | | | | |
| E497 OE2 | 3.3 | N35 ND2 | | | | |

**Supplementary Table 2. List of interfacing residues in the SAM complex, identified by QT Pisa and PyMOL.**

Yellow indicates highly conserved residues (≥60%), red indicates absolutely conserved residues (across 7 species) from sequence alignments. Italics indicates residue conserved across 7 species of Sam50 and *N. gonorrhoeae* BamA.

| Sam35 | Sam37 | Sam50 | | |
|---|---|---|---|---|
| M1* | M1* | M1* | K311 | G425 |
| F27 | W8 | T144 | E315 | L427 |
| L52 | A23 | G147 | G342 | G471 |
| L73 | L52 | N148 | D364 | G473 |
| P97 | P53 | G176 | F366 | V475 |
| P102 | L55 | T177 | G369 | A481 |
| Y156 | I70 | G221 | G370 | R482 |
| L159 | L74 | H236 | R375 | E484 |
| L250 | N137 | R245 | F377 | L485 |
| L313 | Y138 | S255 | G383 | N486 |
| | T142 | G263 | P384 | P490 |
| | P153 | H272 | D389 | G502 |
| | P159 | D277 | G392 | Q504 |
| | L242 | R279 | G393 | G506 |
| | L279 | P285 | P408 | G508 |
| | P288 | G288 | R417 | F511 |
| | | E296 | N423 | L512 |

**Supplementary Table 3. Conserved residues determined from structure-based sequence alignments.**
Asterisk identifies residues that are not visible in the structure.

| *M. thermophila* Sam50 | | | *N. gonorrhoeae* BamA | | |
|---|---|---|---|---|---|
| **Residue 1** | **Residue 2** | **Distance (Å)** | **Residue 1** | **Residue 2** | **Distance (Å)** |
| No Interactions Identified | | | A430 O β1 | T790 OG1 β16 | 3.0 |
| | | | W432 N β1 | L788 O β16 | 3.1 |
| | | | W432 O β1 | L788 N β16 | 3.0 |
| | | | Q434 NE2 L1 | F784 O β16 | 3.0 |

Absolutely conserved Sam50 or BamA residue

Highly conserved residue (≥60% for Sam50, ≥80% for BamA)

*Conserved across 7 species Sam50 and N. gonorrhoeae BamA*

**Supplementary Table 4. Lateral gate interactions in Sam50 and BamA.** Lateral gate interactions in Sam50 detergent structure and BamA (PDB:4K3B), identified using PyMOL.

| ***M. thermophila* Sam50** | | | ***N. gonorrhoeae* BamA** | | |
|---|---|---|---|---|---|
| **Loop 6 Residue** | **Residue 2** | **Distance (Å)** | **Loop 6 Residue** | **Residue 2** | **Distance (Å)** |
| N372 ND2 | S510 O β16 | 2.4 | | | |
| N372 O | L512 N β16 | 3.4 | | | |
| D373 OD1 | R340 NH1 β11 | 3.4 | S658 O | N688 ND2 β12 | 2.9 |
| *R375 NH2* | N402 OD1 β12 | 3.5 | *R660 NH2* | E692 OE2 β12 | 2.4 |
| *R375 NH1* | E484 OE2 β15 | 3.0 | *R660 N* | D713 OD2 β13 | 2.9 |
| | | | *R660 NH1* | D713 OD2 β13 | 3.1 |
| | | | *R660 O* | S751 OG β14 | 2.8 |
| G376 O | R500 N L8 | 3.0 | G661 O | Q782 N L8 | 3.0 |
| | | | G661 N | *Q785 OE1 β16* | 2.4 |
| F377 O | *Q504 NE2 β16* | 3.0 | Y662 O | *Q785 NE2 β16* | 2.5 |
| | | | Y662 OH | Y750 O β14 | 3.4 |
| | | | Y662 OH | K774 O L8 | 3.4 |
| K378 N | E498 O L8 | 3.0 | E663 N | E780 O L8 | 3.3 |

Absolutely conserved Sam50 or BamA residue

Highly conserved residue ($\geq$60% for Sam50, $\geq$80% for BamA)

*Conserved across 7 species Sam50 and N. gonorrhoeae BamA*

**Supplementary Table 5. Loop 6 interactions with the Sam50 and BamA β-barrel.** Loop 6 interactions in Sam50 detergent structure and BamA (PDB:4K3B), identified using PyMOL. Rows in table correspond to conserved interactions of loop 6 in Sam50 and BamA, interaction only in one column indicates it is unique to that protein.

| Protein | MtSam37 | MtTom22 |
|---|---|---|
| **Nr of atoms** | | |
| in interface | 160 (6.3%) | 146 (24.0%) |
| on the surface | 1539 (60.9%) | 498 (81.9%) |
| Total | 2528 (100.0%) | 608 (100.0%) |
| **Buried Surface area ($A^2$)** | 1,424.3 (7.9%) | 1,534.9 (21.2%) |
| **Accessible Surface area ($A^2$)** | 18,048.7 (100%) | 7238.8 (100%) |
| **Solvation Energy (kcal/mol)** | -300.6 | -26.7 |
| Salt bridges in interface | 4 | |
| H-bonds in interface | 10 | |
| **Volume ($A^3$)** | 36,180 | |
| **Buried Surface area in cplx ($A^2$)** | 1,479.6 | |

**Supplementary Table 6. QT-Pisa analysis of molecular interfaces in the high-resolution detergent Sam37-MtTom22 model complex.**

**Supplementary Movie 1. Conformational changes in the lateral gate of Sam50.**
Sam50 (blue) morphing between the partially closed conformation (dimer 1) and the open conformation (dimer 3). β1-β4 rotate outward to open the lateral gate. These conformational changes illustrate the flexibility within the Sam50 β-barrel to accommodate a substrate as it folds. Sam35 (orchid), Sam37 (light green) and the cryoEM densities of the SAM complex in nanodiscs (light yellow) are also shown. The nanodisc clearly outlines the membrane for the complex.
